# Programming Cell-Free Biosensors with DNA Strand Displacement Circuits

**DOI:** 10.1101/2021.03.16.435693

**Authors:** Jaeyoung K. Jung, Khalid K. Alam, Julius B. Lucks

## Abstract

Cell-free biosensors are emerging as powerful platforms for monitoring human and environmental health. Here, we expand the capabilities of biosensors by interfacing their outputs with toehold-mediated strand displacement circuits, a dynamic DNA nanotechnology that enables molecular computation through programmable interactions between nucleic acid strands. We develop design rules for interfacing biosensors with strand displacement circuits, show that these circuits allow fine-tuning of reaction kinetics and faster response times, and demonstrate a circuit that acts like an analog-to-digital converter to create a series of binary outputs that encode the concentration range of the target molecule being detected. We believe this work establishes a pathway to create “smart” diagnostics that use molecular computations to enhance the speed, robustness and utility of biosensors.

## INTRODUCTION

Cell-free biosensing is emerging as a low-cost, easy-to-use and field-deployable diagnostic technology that can be applied to detect a range of contaminants related to human and environmental health [1–3]. At their core, these systems consist of two layers: a sensing layer that includes an RNA or protein-based biosensor and an output layer that includes a reporter construct. By genetically wiring the sensing layer to the output layer, a signal can be generated when the target compound binds to the biosensor and activates the expression of the reporter (**Fig. 1**). Ultimately, reactions can be assembled by embedding the biosensors and reporter constructs within cell-free reaction environments, freeze-dried for easy storage and transportation and rehydrated with a sample of interest at the point-of-need [3, 4]. Using this approach, cell-free biosensors have been created for chemical compounds related to human health such as zinc [5] and quorum sensing molecules produced by pathogenic bacteria [6], drugs such as gammahydroxy-butyrate [7] and water contaminants such as fluoride [3], atrazine [8], antibiotics and heavy metals [9] among others.

**Fig. 1.**
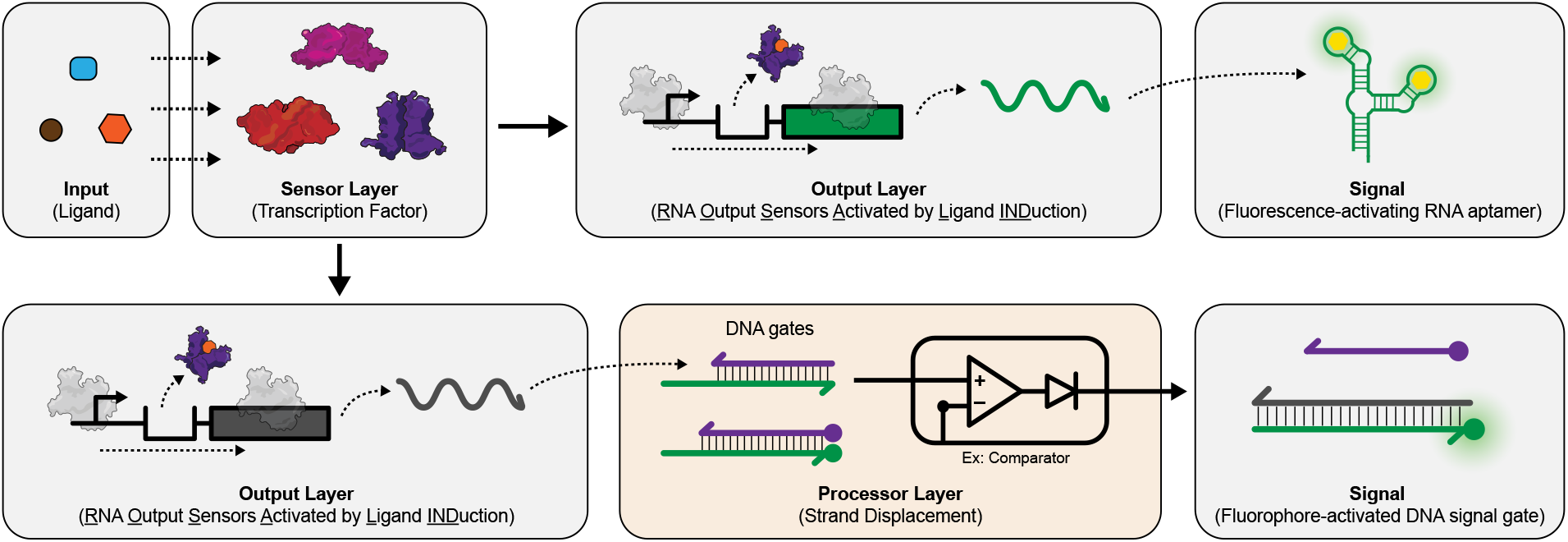
Interfacing cell-free biosensors with a DNA strand displacement circuit information processing layer expands and enhances their function. (Top half) A cell-free biosensor typically activates when a target compound (input) binds to a protein transcription factor (sensor layer) that is configured to activate expression of a reporter construct (output layer). This results in the production of a detectable signal such as fluorescence. (Bottom half) Adding a downstream information processing layer before signal generation can enhance the performance and expand the function of cell-free biosensors by adding computational features such as signal comparison. Here, this is implemented by wiring the biosensing output layer to produce a single stranded RNA capable of activating toehold-mediated strand displacement circuits that generate the signal.

Here, we develop a generalizable strategy to enhance and expand the function of cell-free biosensors by introducing an information processing layer that can manipulate biosensor signals before final signal generation (**Fig. 1**). Such information processing layers are a natural feature of biological organisms and are present in sophisticated genetic networks that enable cells to activate stress responses, alter physiology, guide development and make behavioral decisions based on intracellular and extracellular cues [10]. As such, genetic information processing layers have been extensively leveraged and engineered in synthetic cellular sense-and-respond systems [11, 12]. Similarly, it was recently shown that RNA-based circuits that implement genetic logic and feedback can be added to cell-free biosensing systems to improve their specificity and sensitivity without having to engineer the protein sensors [9]. However, these circuits still directly act on either the sensing or the output layer, limiting the ability to further expand the function of biosensing systems using this approach.

To create a more generalized information processing layer in a cell-free context, we leveraged the development of toehold-mediated DNA strand displacement (TMSD) – a computationally powerful and versatile DNA nanotechnology platform that can be used for information processing *in vitro* [13]. In TMSD, single-stranded DNA (ssDNA) inputs interact with double-stranded DNA (dsDNA) ‘gates’ that are designed to exchange strands and produce ssDNA outputs. By configuring the DNA gates into different network architectures, a range of information processing operations can be performed including signal restoration [14], signal amplification [15] and logic [16, 17], much like a general chemical computational architecture [18]. The well-characterized thermodynamics of DNA base pairing enable large programmable networks to be built from relatively simple building blocks. In addition, the kinetics of these reactions can be precisely tuned by changing the strength of the “toeholds” – single-stranded regions within the DNA gates that initiate the strand displacement process [19]. In this way, TMSD has been used to create a range of devices including *in vitro* oscillators [20], catalytic amplifiers [21], autonomous molecular motors [22, 23] and reprogrammable DNA nanostructures [24, 25]. Furthermore, TMSD circuits capable of sophisticated molecular computations such as complex arithmetic [26] and even molecular neural networks that recognize chemical patterns [27] have been designed. Thus, there is a great potential in utilizing TMSD-based information processing to enhance and expand cell-free biosensor function.

Here, we interface the sensing layers of a previously developed cell-free biosensing platform called ROSALIND [9] with TMSD circuits to expand the platform’s capabilities. It consists of a highly processive phage RNA polymerase (RNAP), an allosteric transcription factor (aTF) and a DNA template that together regulate the synthesis of an invading RNA strand that can activate fluorescence from a DNA signal gate – a dsDNA consisting of a quencher strand and a fluorophore strand with a toehold region. We show that the design of the DNA gate can be optimized to enable T7 RNAP driven *in vitro* transcription (IVT) and TMSD within the same reaction. Next, we systematically develop design principles for optimizing the secondary structure of the synthesized RNA to tune the kinetics of TMSD, notably improving the biosensing response speed. We also apply this principle to interface TMSD with several different aTFs to create biosensors for their cognate ligands. Finally, we address a current limitation of cell-free biosensors by using a model-driven approach to design and build a multi-layer TMSD circuit that acts like an analog-to-digital converter to create a series of binary outputs that encode the concentration range of the target molecule being sensed. Taken together, this work demonstrates that the combination of TMSD and cell-free biosensing reactions can implement molecular computations to enhance the speed and utility of biosensors.

## RESULTS

### Engineering TMSD to be compatible with *in vitro* transcription

To interface cell-free biosensors with TMSD, we first sought to directly interface unregulated IVT reactions with the DNA gates used to generate signals in TMSD. This required us to first validate that a single-stranded RNA can strand-displace a DNA signal gate. The DNA signal gate was created by annealing two ssDNA strands: (1) a fluorophore strand consisting of a 24-nucleotide (nt) ssDNA modified with a 6’ FAM fluorophore on its 5’ end and (2) a quencher strand consisting of a 16-nt ssDNA strand complementary to the fluorophore strand and modified with an Iowa black quencher on its 3’ end (**Supplementary Data File 1**) [28]. Once annealed, the DNA signal gate has an 8-nt toehold on its 3’ end to initiate strand displacement. Following the TMSD design architecture, we designed an invading RNA strand (“InvadeR”) to be fully complementary to the 24-nt fluorophore strand so that it could bind to the toehold region and strand-displace the quencher strand to generate a fluorescent output (**Fig. 2a**).

**Fig. 2.**
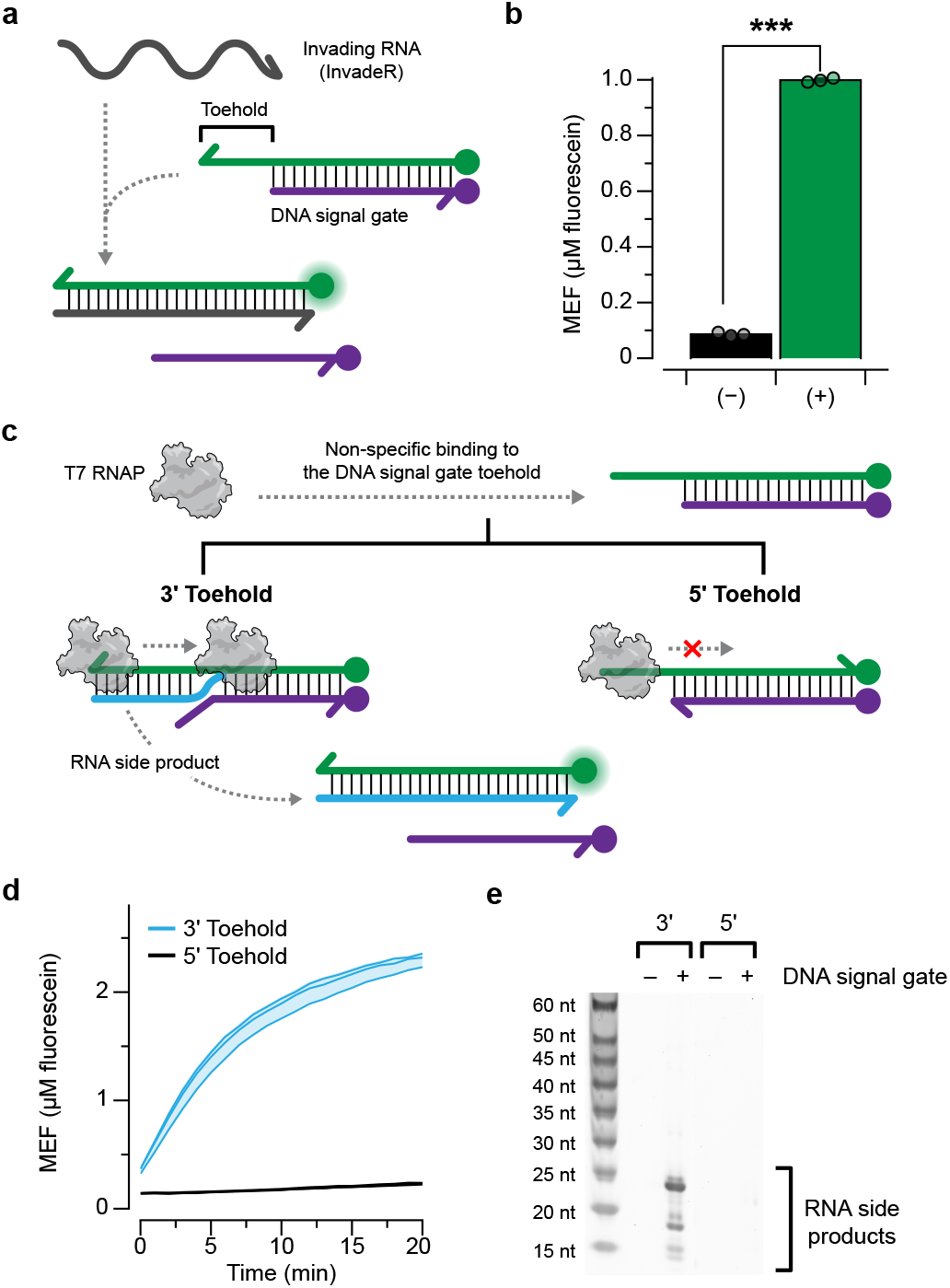
Toehold-mediated DNA strand displacement can be used to track RNA output with an appropriately designed DNA gate. **a,** A DNA signal gate is composed of a 24-nt ssDNA strand modified with a 6’ FAM fluorophore on its 5’ end annealed to a 16-nt ssDNA strand modified with an Iowa black quencher on its 3’ end, leaving an 8-nt toehold on the 3’ end. When an invading RNA strand (InvadeR) complementary to the fluorophore strand is added, it displaces the quencher strand via toehold-mediated strand displacement and activates fluorescence. **b**, When 10 μM of InvadeR is added to an equimolar amount of the DNA signal gate, fluorescence activation is observed. A two-tailed, heteroscedastic Student’s t-test against the no-InvadeR condition was performed, and its P value range is indicated with an asterisk (***P < 0.001, **P = 0.001–0.01, *P = 0.01–0.05). The exact P value along with its degree of freedom can be found in **Supplementary Data File 3**. **c**, In the presence of IVT components, T7 RNAP can non-specifically bind to the toehold region of the DNA signal gate. When the overhanging toehold is on the 3’ end of the gate, this non-specific binding leads to transcription of unwanted RNA side products that can displace the quencher strand. This process is blocked when the overhanging toehold is on the 5’ end of the gate. **d**, The 3’ toehold DNA signal gate leads to fluorescence activation in the presence of T7 RNAP, while the 5’ toehold DNA signal gate does not get activated by T7 RNAP. **e**, When the reaction products from **d** were extracted and run on a polyacrylamide gel, RNA side products appear only when the toehold is located on the 3’ end of the gate. Data shown for n=3 technical replicates as points (**b**) with bar heights representing the average or as lines (**d**) with raw fluorescence standardized to MEF (μM fluorescein). Data shown for n=1 in **e**. Error bars (**b**) and shading (**d**) indicate the average value of the replicates ± standard deviation. The uncropped, unprocessed gel image shown in **e** is available as **Supplementary Data File 2**.

We tested the strand displacement efficiency of this DNA signal gate by adding purified InvadeR to the reaction and monitoring fluorescence, which was standardized to an external fluorescein standard (**Supp. Fig. 1**). Addition of InvadeR resulted in significant fluorescence activation over a no InvadeR control (**Fig. 2b**). In this way, InvadeR behaved similarly to an invading ssDNA strand (“InvadeD”), though these molecules differed in the fluorescence dose response observed (**Supp. Fig. 2a, b**). Specifically, titration of InvadeR resulted in a plateau of fluorescence at lower concentrations than InvadeD. This could be due to differences in RNA and DNA folding thermodynamics. In particular, NUPACK [29] predicts that InvadeD has a less stable structure than InvadeR (–1.43 kcal/mol vs. –3.12 kcal/mol, respectfully) (**Supp. Fig. 2c**), and that InvadeR can bind to itself to form a duplex (**Supp. Fig. 2d, e, Supplementary Data File 2**). This could inhibit binding and strand displacement of the DNA signal gate and lead to a much slower TMSD response even at a higher InvadeR concentration. While this result shows that RNA-initiated TMSD is possible, it also points to several RNA-specific challenges that need to be addressed.

We next sought to determine if InvadeR can be made *in situ* with the DNA signal gate to generate a fluorescent output. Following the ROSALIND platform design, we chose reaction conditions that use the fast phage polymerase, T7 RNAP. We configured the DNA template encoding InvadeR to consist of the minimal 17-base pair (bp) T7 promoter sequence followed by two initiating guanines and the InvadeR sequence. To begin, we tested whether adding T7 RNAP along with other IVT reagents could interfere with the DNA signal gate. To our surprise, we observed an increase in fluorescence in the absence of a T7 DNA transcription template when only T7 RNAP, IVT buffer and NTPs were added to the DNA signal gate (**Fig. 2d**). Previous literature reported the ability of T7 RNAP to initiate promoter-independent transcription from exposed, linear ssDNA regions [30–33]. Based on these observations, we hypothesized that T7 RNAP was initiating transcription from the 3’ end toehold region of the DNA signal gate, causing strand displacement and signal generation (**Fig. 2c**). To test this hypothesis, we reversed the polarity of the DNA signal gate so that the toehold region is on its 5’ end while keeping its sequence the same to prevent T7 RNAP initiation. As expected, no fluorescence signal was observed from the 5’ toehold DNA signal gate in the absence of the DNA template (**Fig. 2d**). To confirm that the signal generation is due to transcription of the DNA signal gate, RNA species from each IVT reaction without the DNA template were extracted and run on a urea-PAGE gel. The resulting gel image shows that RNA side products were generated only from the reaction with the 3’ toehold DNA signal gate (**Fig. 2e, Supplementary Data File 2**). We also designed a modified DNA signal gate where the fluorophore strand is modified with 2’-*O*-methlyation to prevent transcription while keeping the toehold on the 3’ end [32]. As expected, the signal remained at a basal level with the 2’-*O*-methylated DNA signal gate, and no RNA side products were observed on a gel (**Supp. Fig. 3**, **Supplementary Data File 2**), further confirming our hypothesis.

Together, these results revealed several important design features required to interface TMSD with IVT reactions.

### Interfacing *in vitro* transcription with TMSD outputs

We next sought to use TMSD to directly track the RNA outputs generated by T7 RNAP-driven IVT *in situ*. In particular, we focused on optimizing the design of InvadeR for rapid, robust signal generation. Based on our observations in the differences between InvadeR and InvadeD (**Supp. Fig. 2**), we hypothesized that the secondary structure of InvadeR would play a critical role in the strand displacement efficiency and thus the overall TMSD reaction kinetics. For example, an InvadeR strand that forms a stable secondary structure would have to overcome a greater energy barrier to unfold and displace the quencher strand from the DNA signal gate. A stable 3’ end structure of InvadeR could also interfere with the initial binding of the toehold region and hinder the TMSD branch migration process [34, 35].

To test this hypothesis, we designed three different variants of InvadeR that can strand-invade the DNA signal gate optimized in the previous section (**Fig. 3a**). Variant 1 consists of two initiating guanines followed by the sequence fully complementary to the fluorophore strand of the DNA signal gate. For variants 2 and 3, additional nucleotides were inserted between the initiating guanines and the InvadeR sequence to destabilize the predicted G-C base pairs on the 3’ end as well as the overall secondary structure. We also designed strengthened versions of variants 2 and 3 such that the additional nucleotides made the predicted secondary structures more stable with high base pairing probabilities on their 3’ ends. When an equimolar amount of each gel-purified InvadeR variant was added to the DNA signal gate (**Fig. 3b**), we observed fluorescence signals that were ranked according to the predicted minimum free energies of each variant, with variant 3 (–2.8 kcal/mol) showing the highest fluorescence and variant 1 (–5.7 kcal/mol) the lowest (**Fig. 3c**). Furthermore, each strengthened version showed significantly lower fluorescence than the corresponding un-strengthened variant (fold reduction of 2.51 and 2.34 for variants 2 and 3, respectively), confirming our hypothesis (**Fig. 3c**).

**Fig. 3.**
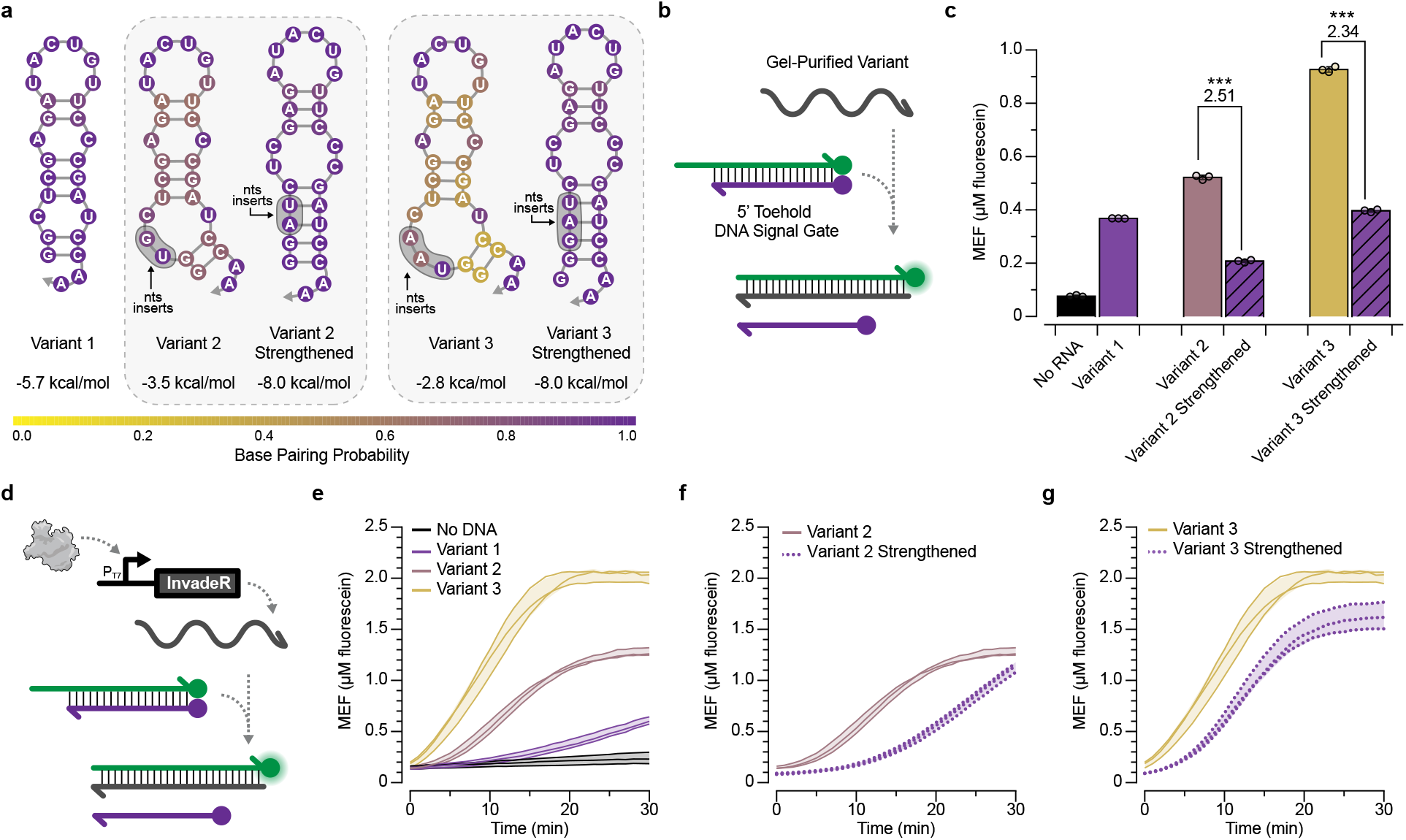
Secondary structure of InvadeR impacts strand displacement efficiency. **a**, Three different variants of InvadeR are designed. Variant 1 includes the two initiating guanines followed by the sequence fully complementary to the fluorophore strand. For variant 2 and 3, two or three additional nucleotides were inserted between the initiating nucleotides and the InvadeR sequence (shaded regions), such that they disrupt the secondary structure at its base. Strengthened versions are created by mutating the additional nucleotides to strengthen the structure, while keeping the number and positions of the nucleotides that interact with the DNA signal gate consistent. Minimum free energies and base pairing probabilities for each structure are predicted using NUPACK at 37° C [29]. **b**, When a gel-purified variant is added to the DNA signal gate, a fluorescent signal via toehold-mediated strand displacement is observed. **c**, 5 μM of gel-purified InvadeR variants were added to an equimolar amount of the DNA signal gate, and fluorescence activation was quantified. Variant 3 generates the highest fluorescent signal followed by variant 2 and 1, while both strengthening mutants show a decrease in signal from their respective variants by the fold reduction indicated above the bars. Two-tailed, heteroscedastic Student’s t-tests were performed between the variants and their respective strengthened versions, and their P value ranges are indicated with asterisks (***P < 0.001, **P = 0.001-0.01, *P = 0.01-0.05). Exact P values along with degrees of freedom can be found in **Supplementary Data File 3**. **d**, When a DNA template encoding InvadeR is included with T7 RNAP and the DNA signal gate, the RNA output can be tracked *in situ* by monitoring fluorescence activation from the signal gate. **e**, Comparison of fluorescence kinetics of the three variants from IVT using an equimolar DNA template (50 nM) or a no template negative control. Comparison of fluorescence kinetics between variants and their strengthening mutants for **f**, variant 2 and **g**, variant 3, shows that strengthening base pairs negatively impact fluorescence kinetics. Data shown for n=3 technical replicates as points (**b**) with bar heights representing the average or n=3 independent experimental replicates as lines (**d-f**) with raw fluorescence standardized to MEF (μM fluorescein). Error bars (**b**) and shading (**d-f**) indicate the average value of the replicates ± standard deviation.

Next, we tested the strand displacement reaction kinetics of the variants transcribed *in situ*. 50 nM of the DNA template encoding each InvadeR variant was added to a reaction mixture containing IVT buffer, T7 RNAP, NTPs and the DNA signal gate, and their fluorescence activation was measured (**Fig. 3d**). We observed fastest fluorescence activation from variant 3 followed by variants 2 and 1 (**Fig. 3e**), which agree with the previous experiment. When we compared the reaction kinetics of variant 2 and 3 to that of their respective strengthened versions, we observed slower responses from the strengthened versions, reaffirming our hypothesis (**Fig. 3f, g**).

However, we observed some discrepancies between the predicted secondary structures and the reaction kinetics of the strengthened variants. For instance, while NUPACK predicts lower minimum free energy values from the strengthened variants than variant 1, the strengthened variants show faster reaction kinetics than variant 1 (**Fig 3e–g**). Furthermore, the endpoint fluorescence values reached by the strengthened variants *in situ* are higher than that of variant 1, conflicting with the results observed in **Fig. 3c**. We hypothesized that these discrepancies could be due to varying transcription efficiency of each DNA template, as it has been reported that the sequence of the initially transcribed region greatly impacts the T7 RNAP transcription efficiency [36]. To test this hypothesis, RNA products from 30-minute long IVT reactions initiated with each DNA template were extracted, and their RNA concentrations were measured using both the RNA Qubit assay and a gel band intensity analysis from a urea-PAGE gel stained with SYBR gold (**Supp. Fig. 4**) [37]. In both cases, we observed the highest RNA concentrations from the strengthened variants (**Supp. Fig. 4b, e**). Despite having the lowest transcription efficiency, variant 3 still showed the fastest kinetics, indicating that the secondary structure greatly affects the TMSD response speed. We also found that adding a T7 terminator sequence at the end of the DNA template speeds up the reaction [38], although not considerably (**Supp. Fig. 5**).

Together, these results show that both secondary structure and transcription efficiency impact the ability of RNA strands to invade DNA signal gates and that these design principles can be leveraged to enhance reaction speed.

### Interfacing cell-free biosensors with TMSD outputs

Next, we sought to determine whether the transcription of InvadeR can be regulated with an aTF, thus creating a ligand-responsive biosensor that uses TMSD outputs. This required us to insert an aTF operator sequence in between the T7 promotor and InvadeR sequence to allow an aTF to regulate transcription. We previously demonstrated that the spacing between the minimal 17-bp T7 promoter sequence and the aTF operator sequence is important for efficient regulation of IVT in ROSALIND reactions [9]. To test if this spacing remained important in the TMSD platform, we used TetR as our model aTF to determine the optimum spacing for efficient repression in the presence of TetR and efficient transcription in the absence of TetR [39]. We used the native sequence that follows the canonical T7 RNAP promoter as a spacer in 2-bp increments from 0 to 10-bp, immediately followed by the *tetO* operator sequence and InvadeR sequence (**Fig. 4a**). IVT reactions were set up using 50 nM DNA template with or without purified recombinant TetR protein in 100-fold excess of the DNA template. Consistent with our previously reported observation [9], the absence of any spacing resulted in no fluorescence activation in either the presence (regulated) or the absence (unregulated) of TetR (**Fig. 4b**). However, robust fluorescence signal was observed without TetR when using a 2-bp spacer, which was reduced to nearly baseline levels when regulated by TetR. Spacers longer than 2-bp resulted in T7 RNAP read-through, leading to fluorescence activation in the presence of TetR. We note, however, if an operator sequence starts with a guanine, thus acting as the initiating nucleotide for T7 RNAP, no spacing could still lead to transcription in the absence of the corresponding aTF.

**Fig. 4.**
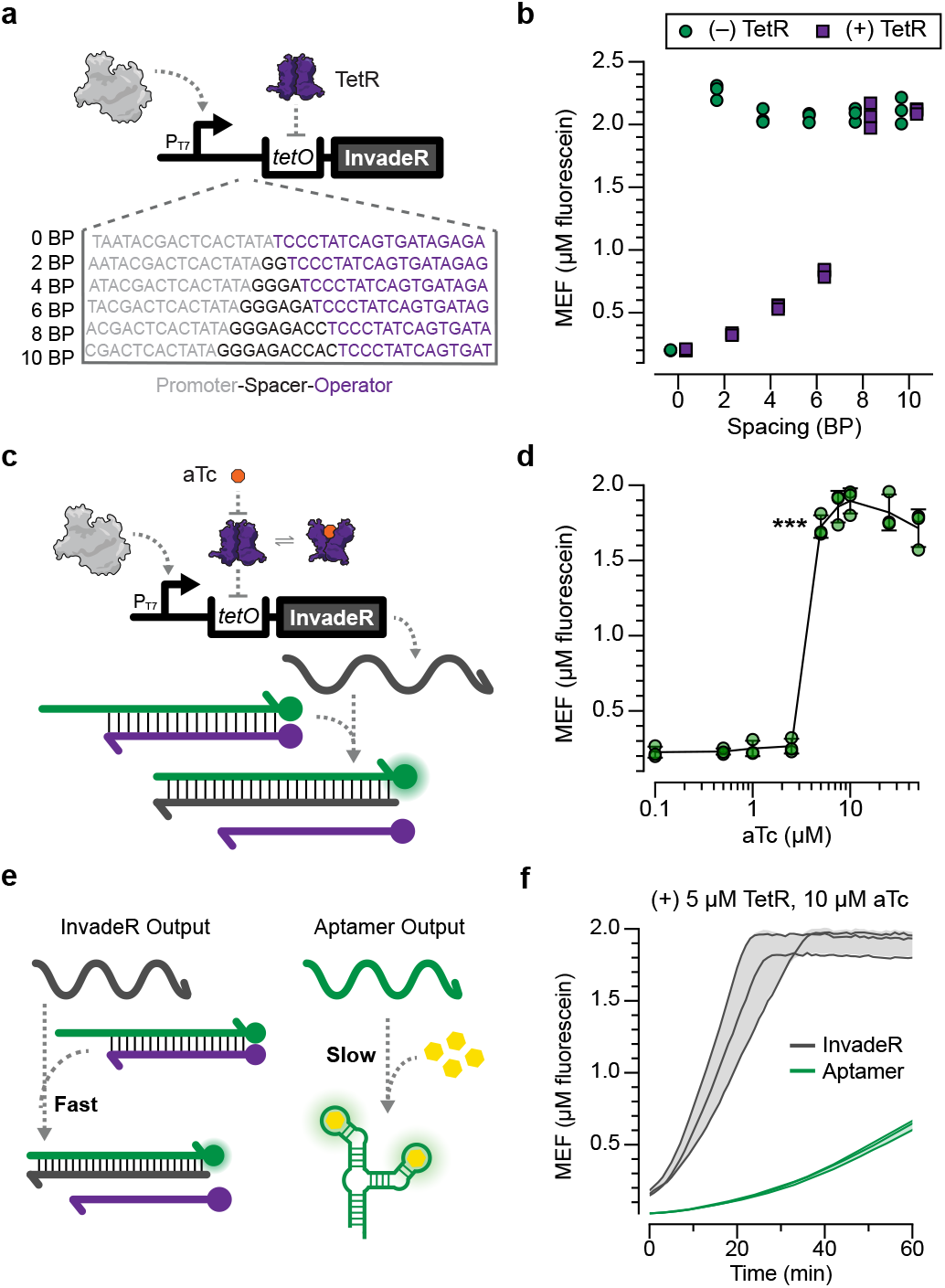
Transcription of InvadeR can be regulated with an allosteric transcription factor. **a**, IVTs can be allosterically regulated with a template configured to bind a purified transcription factor (TetR) via operator sequence (*tetO*) placed downstream of the T7 promoter. A series of spacers in 2-bp intervals was constructed to evaluate the impact of spacer length on the ability of TetR to regulate the transcription of InvadeR. **b**, End-point data (at 1h) shown for promoter-operator spacer variants regulated (with 5 μM TetR dimer, 50 nM DNA template) and unregulated (without TetR). **c**, Induction of a TetR-regulated IVT reaction occurs in the presence of the cognate ligand, anhydrotetracycline (aTc), which binds to TetR and prevents its binding to *tetO*. This allows transcription to proceed, leading to fluorescence activation via toehold-mediated strand displacement. **d**, Dose response with aTc, measured at 1h with 50 nM DNA template and 5 μM TetR dimer. The lowest ligand concentration at which the signal is distinguishable from the background was determined using a two-tailed, heteroscedastic Student’s t-test against the no-ligand condition, and its P value range is indicated with an asterisk (***P < 0.001, **P = 0.001-0.01, *P = 0.01-0.05). Exact P values along with degrees of freedom for all ligand concentrations tested can be found in **Supplementary Data File 3**. Data for no-ligand condition were excluded because the x-axis is on the log scale and are presented in **Supplementary Data File 3. e**, The speed of the toehold-mediated strand displacement output is fast and tunable while that of the RNA aptamer output is slow and difficult to alter. **f**, Comparison of fluorescence kinetics between the TetR-regulated InvadeR and the aptamer outputs when induced with 10 μM aTc. All data shown for n=3 independent experimental replicates as points (**b, d**) or lines (**f**) with raw fluorescence values standardized to MEF (μM fluorescein). Error bars (**d**) and shading (**f**) indicate the average value of the replicates ± standard deviation.

Using the 2-bp spacer, we next determined whether TetR can be de-repressed with its cognate ligand, anhydrotetracycline (aTc) to allow transcription of InvadeR (**Fig. 4c**). When a range of aTc concentrations was added to reactions each containing T7 RNAP, 50 nM DNA template, 5 μM TetR dimer and 5 μM DNA signal gate, we observed a strong repression over 1h down to low micromolar amounts of aTc, with half-maximal induction between 2.5 μM and 5 μM of aTc. (**Fig. 4d**).

Due to the rapid speed of TMSD reactions [19], we hypothesized that the ligand-mediated induction speed of the InvadeR output would be much faster than the previously used fluorescence-activating RNA aptamer output especially because the fluorescence activation from the aptamer is limited by the kinetics of dye binding (**Fig. 4e**). Specifically, the kinetic rate of TMSD of an 8-nt toehold is two orders of magnitude higher than the Spinach aptamer and DFHBI-1T binding kinetic rate [34, 40]. Furthermore, it has been recently discovered that fluorescence-activating RNA aptamers are prone to misfolding [41], possibly contributing to slower and reduced signal generation relative to the amount of RNA transcribed. As expected, we observed that the InvadeR platform activates fluorescence visible in ~10 minutes which is approximately 5-times faster than the RNA aptamer platform when using the equimolar amounts of the DNA template, TetR and aTc (**Fig. 4f**).

Overall, these results demonstrate that an aTF-based biosensor can be successfully interfaced with TMSD outputs, leading to immediate improvements in reaction speed.

### Optimizing invading RNA designs for different biosensor families

Having demonstrated the ability to regulate InvadeR with TetR, we next sought to determine whether the system is compatible with different families of aTFs to create biosensors for different classes of chemical contaminants. In addition to TetR, we chose TtgR [42] and SmtB [43] as representative aTFs of the MarR family [44] and SmtB/ArsR family [45], respectively. We placed the cognate operator sequence of each aTF 2-bp downstream of the T7 promoter and immediately upstream of the InvadeR sequences (**Fig. 5a–c**). When tetracycline was used to induce TetR-regulated reactions, we observed a strong and robust fluorescent signal visible in ~10 minutes (**Fig. 5d, Supp. Fig. 6a**). Similarly, when a cognate ligand of TtgR, naringenin was added to TtgR-regulated reactions, we again saw robust fluorescence activation only in the presence of the ligand (**Fig. 5e, Supp. Fig. 6b**). We note that we observed a slight decrease in the induction speed from the TtgR-regulated reactions compared to that of the TetR-regulated reactions, possibly due to the reduced *ttgO* DNA template concentration needed to accommodate TtgR’s relatively weak binding affinity to its operator sequence [46].

**Fig. 5.**
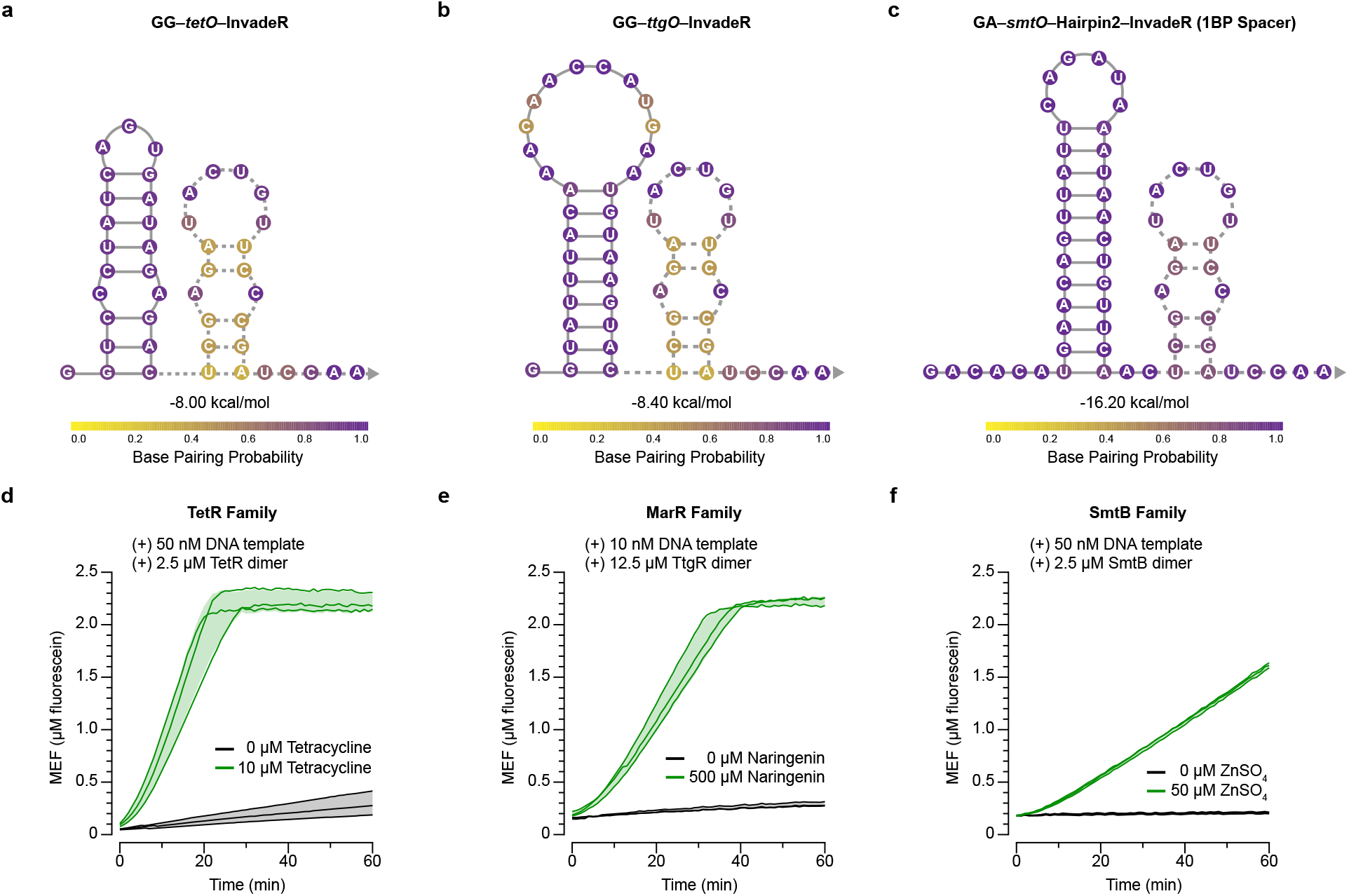
InvadeR can be modularly configured with an operator sequence to sense various molecules. DNA templates encoding InvadeR are modified to contain the T7 promoter followed by a 2-bp spacer and an aTF operator sequence immediately upstream of the InvadeR sequence. Secondary structures, minimum free energies and base pairing probabilities of **a**, *GG-tetO-* InvadeR, **b**, GG-*ttgO*-InvadeR and **c**, GA-*smtO*-Hairpin2-InvadeR (1BP spacer variant) are predicted using NUPACK at 37° C [29]. The GA-*smtO*-Hairpin2-InvadeR sequence includes an RNA hairpin designed to minimize structural interference of InvadeR with *smtO* and optimize the signal (**Supp. Fig. 7h, i**). The sequence complementary to the fluorophore strand is denoted with a dotted line. **d**, TetR can be used to sense tetracycline. **e,** TtgR, a MarR-family aTF, can be used to sense naringenin. **f**, SmtB can be used to sense zinc. Variations in activation kinetics match the trend where the predicted secondary structure of the modified InvadeR impacts the induction speed. All data shown for n=3 independent experimental replicates as lines with raw fluorescence value standardized to MEF (μM fluorescein). Shading indicates the average value of the replicates ± standard deviation.

We were immediately successful in adapting the system to TetR and TtgR, which we hypothesize was due to the fact that introducing *tetO* and *ttgO* minimally impacted the secondary structure of InvadeR since each RNA-encoded operator sequence was predicted to form a hairpin within itself (**Fig. 5a, b**). In contrast, introducing the *smtO* sequence resulted in much slower reaction speeds, even in unregulated reactions (**Supp. Fig. 7c**). Interestingly, we observed that the *smtO* sequence is predicted to form a strong hairpin with the InvadeR sequence (**Supp. Fig. 7a**), which based on our above results could impact reaction speeds. We hypothesized that adding extra sequences between the *smtO* and InvadeR sequences to prevent intramolecular *smtO*:InvadeR folding would improve the reaction kinetics. To test this hypothesis, we first concatenated an additional InvadeR sequence in series so that the first InvadeR sequence forms a hairpin with *smtO,* leaving the second InvadeR sequence available to strand-displace the DNA signal gate (**Supp. Fig. 7b**). As expected, we observed a pronounced improvement in the response speed (**Supp. Fig. 7c**).

However, it is likely that the speed improvement is due to not only the structural change of the transcript, but also a single transcript potentially strand-invading two DNA signal gates. To decouple this effect, we designed two different *smtO*-hairpin-InvadeR variants where we used NUPACK to design a sequence that would form a hairpin with the *smtO* sequence (**Supp. Fig. 7d**, **e**). We observed varying degrees of speed improvement from these variants (**Supp. Fig. 7f**). To further investigate different design features of these variants, we took the *smtO*-hairpin2-InvadeR sequence and designed a series of additional variants either by lengthening the predicted stem-loop of the hairpin or the spacer sequence between the hairpin and InvadeR (**Supp. Fig. 7g**). We observed that the response speed increases when decreasing the stem-loop length (**Supp. Fig. 7h**) and increasing the spacer length (**Supp. Fig. 7i**). Using this combination of features, we created an improved design that showed clear activation of a SmtB-regulated reaction in the presence of ZnSO_4_ with low background signal (**Fig. 5c, f**).

Together, these results demonstrate that the modularity of the ROSALIND platform is extensible to the TMSD platform. These results also reinforce that the secondary structures of the invading RNA strands play a critical role in determining reaction speed and that these structures can be computationally designed to improve the TMSD response speed.

### Using a TMSD circuit processing layer to quantify biosensor outputs

The interface of biosensing with TMSD creates a potentially powerful molecular computation paradigm for engineering molecular devices that can perform programmed tasks in response to specific chemical triggers. This is especially true since TMSD circuits are much easier to program than protein-based circuits as a result of their simpler design rules [47], computational models that accurately predict their behavior [34, 35] and the emerging suite of design tools to build TMSD circuits [26, 48]. We, therefore, sought to leverage these features of TMSD circuits to create an information processing layer for cell-free biosensors that could be used to expand their function.

As a model example, we chose to focus on quantifying biosensor outputs. In typical cell-free biosensing systems, the sensor layer is wired to the output layer (**Fig. 1**), thus directly coupling the amount of output signal to the properties of the biosensor. In many cases, this results in an output signal that is generated above a specific detection threshold which is determined by the aTF-ligand and aTF-DNA binding constants [49], making it difficult to obtain information about the input target compound concentration. One approach to solving this challenge is to configure reactions to generate a proportional output response [5], though this system requires users to judge output intensity or hue to estimate the input target concentration which can lead to uncertainty. Alternatively, we chose to create a system similar to an analog-to-digital converter (ADC) circuit – widespread in electronic systems that interface sensors to information processing modules [50] – that creates a series of binary outputs that encode the analog input concentration of the target compound (**Supp. Fig. 8**).

To construct a genetic ADC circuit, we first needed to create a comparator circuit – a building block of ADCs that produces a “True” binary output when the input is above a pre-defined threshold. ADC circuits can then be built by creating a series of comparators, each with different thresholds. Previously, this concept of thresholding was implemented in *in vitro* DNA-only TMSD circuits to act as a low-level noise filter [14, 26]. Thresholding can be implemented in TMSD because the reaction kinetics of strand displacement can be precisely tuned by adjusting the length of DNA gate toehold regions [19]. Specifically, each additional nt added to a toehold region enhances strand displacement kinetics by 10-fold [34]. As a result, additional DNA gates with longer toeholds can be designed to preferentially react with inputs, thus only allowing DNA signal gates to be activated when the input DNA strand completely consumes the longer-toehold DNA gates.

Our first step was to build a similar thresholding circuit but using input RNA strands generated *in situ*. The DNA threshold gate was designed to contain two strands: an identical strand to the fluorophore strand of the signal gate and a shortened complementary strand to allow a longer 8-nt toehold compared to the 4-nt toehold of the signal gate (**Fig. 6a**). Additionally, the threshold gate lacked the fluorophore and quencher modifications. In this design, InvadeR should react preferentially with the threshold gate with orders of magnitude increased rates, preventing InvadeR from interacting with the signal gate. Only after the threshold gate is completely exhausted can InvadeR efficiently strand-invade the signal gate to generate a fluorescent signal. We reasoned that by tuning the amount of the threshold gate present in the reaction, we can precisely control the time at which InvadeR activates fluorescence from the signal gate. Modeling this kinetic behavior using a set of ordinary differential equations (ODEs) that describe the reactions (IVT and TMSD) in the system (see **Supplementary Method** for details on the ODE model used) showed that this is indeed the predicted behavior of the setup (**Fig. 6b**). We then tested these reactions experimentally and observed quantitative agreement with the model predictions (**Fig. 6b**). In this way, a thresholded TMSD reaction acts as a “kinetic” comparator circuit – for a given input, the time at which signal generation occurs is proportional to the amount of the threshold gate added to the reaction.

**Fig. 6.**
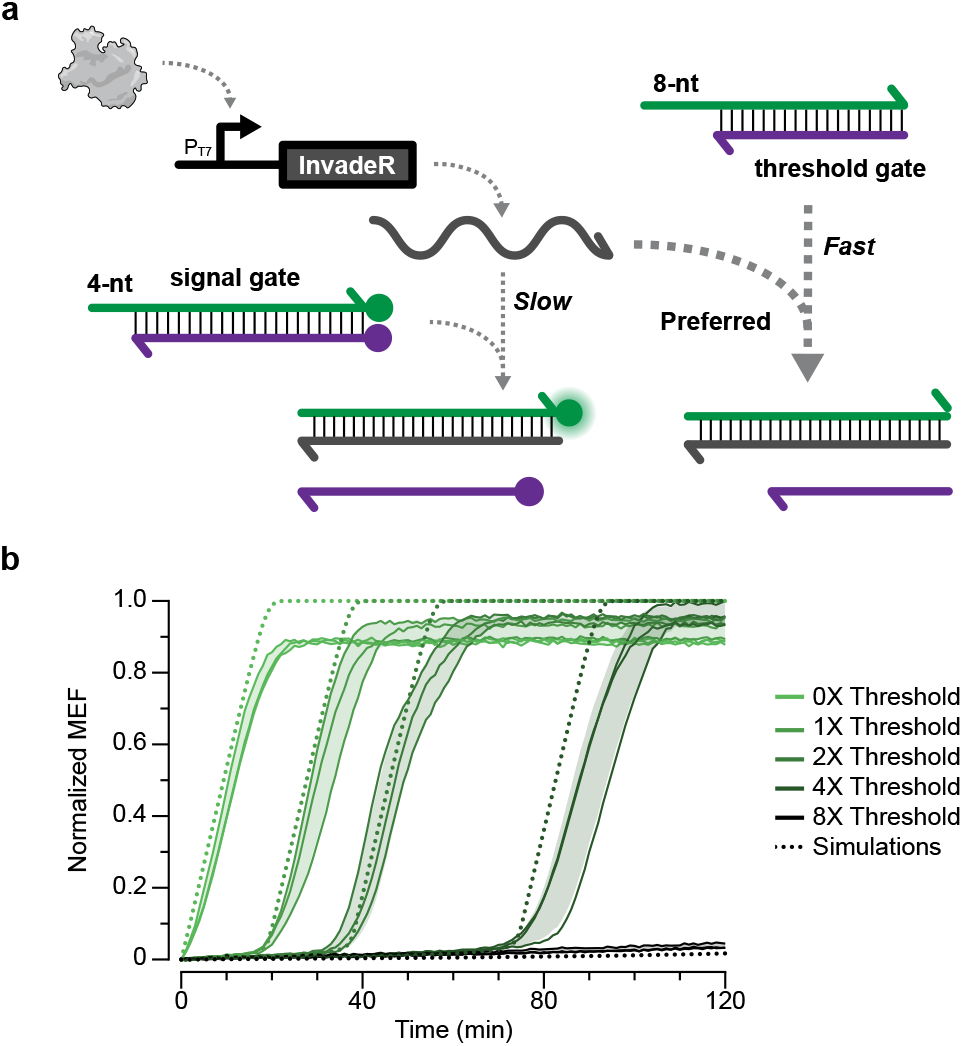
An additional DNA gate can be applied to quantitatively sequester the RNA output. **a,** Increasing the length of the DNA gate toehold region can be used to speed the strand invasion process. An unlabeled DNA gate with a longer toehold (8-nt) can then preferentially react with InvadeR, acting as a programmable threshold. InvadeR can only strand-displace the signal gate (4-nt toehold) after the threshold gate is exhausted. **b**, Titrating the 8-nt toehold threshold gate in different ratios above a fixed signal gate concentration (0X-8X) results in a time delay in fluorescence activation that can be quantitatively modeled with ODE simulations (dotted lines). All data shown for n=3 independent experimental replicates as lines. Raw fluorescence values were first standardized to MEF (μM fluorescein) and normalized to the maximum MEF among all conditions to accommodate their comparison to the simulations (See **Materials and Methods** for the normalization method used). Shading indicates the average value of the replicates ± standard deviation.

Next, we sought to create a series of biosensing TMSD comparator circuits to act as an ADC for ligand concentration. Specifically, we prepared a strip of reactions where each tube contains a different amount of the threshold gate. By adding the same input ligand concentration to each tube and observing the output at a specific time point, a user can count the number of activated tubes to obtain semi-quantitative information about ligand concentration (**Fig. 7a**). For example, since a higher ligand concentration is required to overcome a higher threshold value, we expected to observe different numbers of activated tubes depending on the input concentration – the higher the input, the greater the number of activated tubes.

**Fig. 7.**
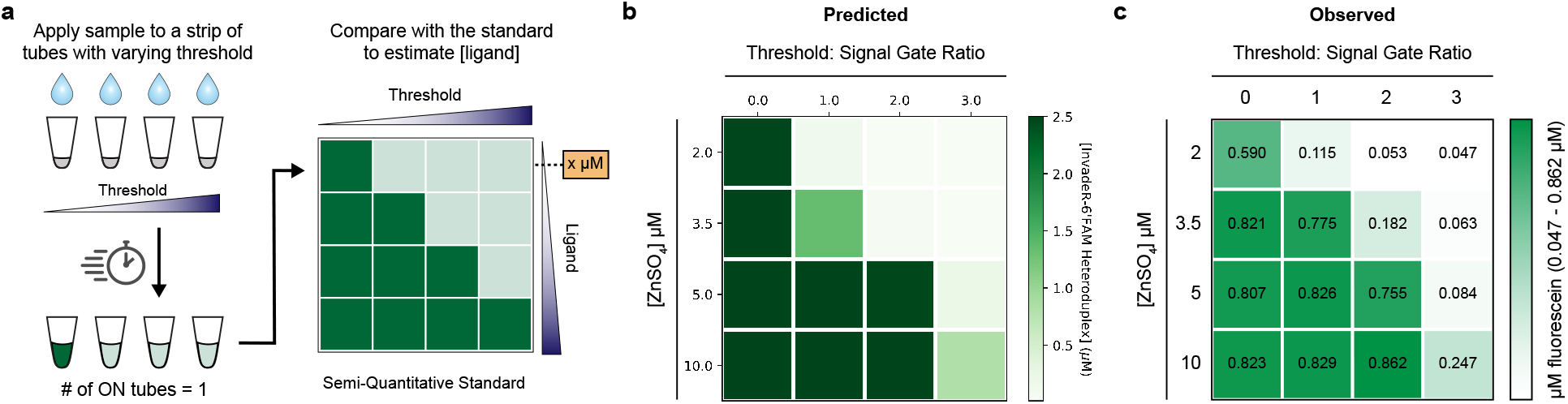
Quantifying ligand concentration with a genetic ADC circuit. **a,** A genetic analog-to-digital (ADC) circuit is made by constructing a strip of tests of the same sensor, with each test containing a different concentration of the DNA threshold gate. A higher threshold gate concentration requires a higher ligand concentration to activate fluorescence. When the same sample is applied to each tube, a user can obtain semi-quantitative information about the concentration of ligand present in the sample (analog input) by counting the number of tubes that activate (binary digital output). Characterization of a genetic ADC circuit for zinc using **b,** ODE simulations and **c,** end-point experimental data at 100 min. The values on the heatmap represent the average MEF (μM fluorescein) of n=3 independent experimental replicates (See **Supp. Fig. 9** for all data).

We first built a model for the system to determine the feasibility of the approach using the same set of ODEs used in **Fig. 6b** with the addition of aTF–DNA and aTF–ligand binding kinetics, focusing on zinc sensing with SmtB because of its relevance in municipal water supplies [51]. We used simulations to determine the threshold gate concentrations needed to activate one, two, three or four tubes after 100 minutes corresponding to zinc concentrations of 2 μM, 3.5 μM, 5 μM and 10 μM, respectively (**Fig. 7b**). We then proceeded to build this genetic ADC circuit with the SmtB-regulated TMSD reactions. Four ADC reaction sets were built using the threshold gate concentrations simulated in **Fig. 7b**, and each set was tested with either 2 μM, 3.5 μM, 5 μM or 10 μM ZnSO_4_. When the reactions were run for 100 minutes, we saw the expected pattern of signals where the number of activated tubes increased with higher input ZnSO_4_ concentrations (**Fig. 7c, Supp. Fig. 9a–d**). This implementation allows a user to determine the unknown input zinc concentration range by directly reading out the number of activated tubes.

While simple, this demonstration represents the potential of TMSD circuits as an information processing layer to expand the functionality of cell-free biosensors where the circuits transform an analog input signal into a digital readout to increase ease of interpretation and information content of the output signals.

## DISCUSSION

In this study, we show that nucleic acid strand displacement circuits can be interfaced with IVT to act as an information processing layer for cell-free biosensors. We found that the speed of DNA strand displacement outputs led to a significant enhancement of output signal generation speed, with visible outputs being produced in ~10 minutes compared to ~50 minutes for fluorescent RNA aptamer outputs (**Fig. 4f**). More significantly, we found that the simple and defined nature of ROSALIND, combined with the computational power of TMSD and the ability to accurately model TMSD reactions with ODE simulations, enabled us to layer multiple DNA gates to design and validate a circuit that can estimate the concentration range of an unknown target compound within a sample (**Fig. 7**). This platform is also amenable to lyophilization, although improvements are needed to ensure long-term storage without loss of signal over time (**Supp. Fig. 10**).

While simple in concept, we found that the combination of TMSD with cell-free biosensing reactions did not work immediately. This was due to the incompatibility of 3’ toehold overhangs in DNA gates with T7 RNAP-driven IVT reactions [48]. A careful analysis of the issues determined that this incompatibility is due to undesired transcription of these 3’ toehold overhangs by T7 RNAP, which can be solved by changing toehold overhangs to be on the 5’ ends (**Fig. 2**), or by modifying the DNA gates by incorporating 2’-*O*-methylated nucleotides (**Supp. Fig. 3**) [32].

A key advantage of this platform is its flexibility in designing the DNA gates and invading RNA strands once the basic design principles of the RNA-DNA interactions of this system are understood. In particular, we discovered that intramolecular RNA structures — caused by sequence constraints of the corresponding DNA signal gates or by the incorporation of aTF operator sites — can inhibit the basic strand displacement reactions. We found that nucleic acid design tools such as NUPACK can be used to manipulate or add sequence regions predicted to minimize these structures and therefore enhance compatibility with TMSD. Using this approach, we were able to freely design different InvadeR strands with minimum secondary structures on their toehold binding regions to improve the kinetics for both unregulated (**Fig. 3**) and aTF-regulated reactions (**Supp. Fig. 7**). Furthermore, T7 transcription efficiency can also be altered by optimizing the initially transcribed sequences of an InvadeR strand [36], and this can be leveraged to tune the response speed as well.

One of the major limitations of the platform is its cost. Despite the significant decrease in cost of DNA synthesis, chemically modified oligos with purification can still cost ~$100 USD or more, though a single batch can be used to make hundreds of reactions. Furthermore, DNA gates often need to be gel-purified after hybridization to eliminate any unbound ssDNA strands, which can be time-intensive and laborious. This challenge can be partially solved by designing DNA signal gate sequences to minimize fluorophore quenching by the base adjacent to the modification [52]. Additionally, invading RNA strands can be designed to minimize intra- and intermolecular interactions to ensure that all TMSD reactions go to completion to maximize a fluorescent signal from the amount of a DNA signal gate used.

The key feature of this study was demonstrating the potential of TMSD circuits to expand the function of cell-free biosensors by acting as additional information processing layers. While a similar approach was recently developed to interface aTF-based biosensing with TMSD through endonuclease-mediated TMSD cascades [53], no programmable molecular computation beyond simple contaminant detection was presented. As in natural organisms, information processing layers significantly expand the function of cell-free sensors by enabling systems to manipulate output signals, perform logic operations and make decisions. As a demonstration, we developed a genetic ADC circuit that can be used to estimate an input ligand concentration at a semi-quantitative level (**Fig. 7**). In particular, this genetic ADC circuit uses thresholding computation to convert an analog signal of an input target molecule concentration into a digital output of the number of activated tubes. A key feature of TMSD that enabled this development is its ability to precisely tune reaction rates based on the toehold length. We note, however, this genetic ADC circuit is different from an electrical ADC circuit in that its result depends on time of activation because the circuit relies on thresholding reaction kinetics rather than strictly input concentrations. As a result, this ADC strategy is best suited to distinguishing between ligand concentrations that cause differences in output kinetics. (**Supp. Fig. 9e–g**).

We believe that this platform opens the door to enabling other types of molecular computation in cell-free systems. For example, an amplification circuit such as a catalytic hairpin assembly [54] could be applied to ROSALIND with TMSD for amplifying signals and making a sensor ultrasensitive. Beyond thresholding, other operations demonstrated in DNA seesaw gate architectures could be ported to this platform for various computations [26]. For instance, logic gate operations could be implemented to enable simultaneous detection of multiple ligands or to develop a general strategy to fix aTF ligand promiscuity [9]. In addition, since virtually any aTF can be used, as previously demonstrated in the RNA aptamer platform [9], multiple DNA gates with different reporters could be added for multiplexing. The fundamental role that ADC circuits play in interfacing analog and digital electronic circuitry also holds promise for adopting additional electronic circuit designs to biochemical reactions.

Together, these results show that establishing an interface between biosensing and TMSD circuits is a promising first step towards creating a general molecular computation platform to enhance and expand the function of cell-free biosensing technologies.

## MATERIALS AND METHODS

### Strains and growth medium

*E. coli* strain K12 (NEB Turbo Competent *E. coli*, New England Biolabs #C2984) was used for routine cloning. *E. coli* strain Rosetta 2(DE3)pLysS (Novagen #71401) was used for recombinant protein expression. Luria Broth supplemented with the appropriate antibiotic(s) (100 μg/mL carbenicillin, 100 μg/mL kanamycin and/or 34 μg/mL chloramphenicol) was used as the growth media.

### DNA gate preparation

DNA signal gates used in this study were synthesized by Integrated DNA technologies as modified oligos. They were generated by denaturing a 6-FAM (fluorescein) modified oligonucleotide and the complementary Iowa Black^®^ FQ quencher modified oligonucleotide (**Supplementary Data File 1**) at 95° C separately for 3 minutes and slow cooling (−0.1° C/s) to room temperature in annealing buffer (100 mM potassium acetate and 30 mM HEPES, pH 8.0). Annealed oligonucleotides where then purified by resolving them on 20% native PAGE-TBE gels, isolating the band of expected size and eluting at 4° C overnight in annealing buffer. The eluted DNA gate was then ethanol precipitated, resuspended in MilliQ ultrapure H_2_O and concentration quantified using the Thermo Scientific™ NanoDrop™ One Microvolume UV-Vis spectrophotometer. The DNA threshold gate used in **Fig. 6** and **7** was prepared using the same method but by annealing two complementary oligonucleotides without any modifications.

### Plasmids and genetic parts assembly

DNA oligonucleotides for cloning and sequencing were synthesized by Integrated DNA Technologies. Genes encoding aTFs were synthesized either as gBlocks (Integrated DNA Technologies) or gene fragments (Twist Bioscience). Protein expression plasmids were cloned using Gibson Assembly (NEB Gibson Assembly Master Mix, New England Biolabs #E2611) into a pET-28c plasmid backbone and were designed to overexpress recombinant proteins as C-terminus His-tagged fusions. A construct for expressing SmtB additionally incorporated a recognition sequence for cleavage and removal of the His-tag using TEV protease. Gibson assembled constructs were transformed into NEB Turbo cells, and isolated colonies were purified for plasmid DNA (QIAprep Spin Miniprep Kit, Qiagen #27106). Plasmid sequences were verified with Sanger DNA sequencing (Quintara Biosciences) using the primers listed in **Supplementary Data File 1**.

All transcription templates except for the templates encoding variant 1 in **Fig. 3** and InvadeR in **Fig. 6** were generated by PCR amplification (Phusion High-Fidelity PCR Kit, New England Biolabs #E0553) of an oligo that includes a T7 promoter, an optional aTF operator site, the InvadeR coding sequence and an optional T7 terminator using the primer sets listed in **Supplementary Data File 1**. Here, we define the T7 promoter as a minimal 17-bp sequence (TAATACGACTCACTATA) excluding the first G that is transcribed. Amplified templates were then purified (QIAquick PCR purification kit, Qiagen #28106), verified for the presence of a single DNA band of expected size on a 2% TAE-Agarose gel, and concentrations were determined using the Qubit dsDNA BR Assay Kit (Invitrogen #Q32853). The templates encoding variant 1 in **Fig. 3** and InvadeR in **Fig. 6** were generated using the same method described in **DNA gate preparation** but with two complementary oligonucleotides that include a T7 promoter and the InvadeR coding sequence.

All plasmids and DNA templates were stored at 4° C until usage. A spreadsheet listing the sequences and Addgene accession numbers of all plasmids and oligos generated in this study are listed in **Supplementary Data File 1**.

### RNA expression and purification

InvadeR variants used for the purified oligo binding assays were first expressed by an overnight IVT at 37° C from a transcription template encoding a cis-cleaving Hepatitis D ribozyme on the 3’ end of the InvadeR sequence with the following components: IVT buffer (40 mM Tris-HCl pH 8, 8 mM MgCl_2_, 10 mM DTT, 20 mM NaCl, and 2 mM spermidine), 11.4 mM NTPs pH 7.5, 0.3U thermostable inorganic pyrophosphatase (#M0296S, New England Biolabs), 100 nM transcription template, 50 ng of T7 RNAP and MilliQ ultrapure H_2_O to a total volume of 500 μL. The overnight IVT reactions were then ethanol-precipitated and purified by resolving them on a 20% urea-PAGE-TBE gel, isolating the band of expected size (26 - 29 nt) and eluting at 4° C overnight in MilliQ ultrapure H_2_O. The eluted InvadeR variants were ethanol precipitated, resuspended in MilliQ ultrapure H_2_O, quantified using the Qubit RNA BR Assay Kit (Invitrogen #Q10211) and stored at −20° C until usage. The Hepatitis D ribozyme sequence used can be found in **Supplementary Data File 1**.

### aTF expression and purification

aTFs were expressed and purified as previously described [9]. Briefly, sequence-verified pET-28c plasmids were transformed into the Rosetta 2(DE3)pLysS *E. coli* strain. 1~2 L of cell cultures were grown in Luria Broth at 37° C, induced with 0.5 mM of IPTG at an optical density (600 nm) of ~0.5 and grown for 4 additional hours at 37° C. Cultures were then pelleted by centrifugation and were either stored at −80° C or resuspended in lysis buffer (10 mM Tris-HCl pH 8, 500 mM NaCl, 1 mM TCEP, and protease inhibitor (cOmplete EDTA-free Protease Inhibitor Cocktail, Roche)) for purification. Resuspended cells were then lysed on ice through ultrasonication, and insoluble materials were removed by centrifugation. Clarified supernatant containing TetR was then purified using His-tag affinity chromatography with a Ni-NTA column (HisTrap FF 5mL column, GE Healthcare Life Sciences) followed by size exclusion chromatography (Superdex HiLoad 26/600 200 pg column, GE Healthcare Life Sciences) using an AKTAxpress fast protein liquid chromatography (FPLC) system. Clarified supernatants containing TtgR and SmtB were purified using His-tag affinity chromatography with a gravity column charged with Ni-NTA Agarose (Qiagen #30210). The eluted fractions from the FPLC (for TetR) or from the gravity column (for TtgR and SmtB) were concentrated and buffer exchanged (25 mM Tris-HCl, 100 mM NaCl, 1mM TCEP, 50% glycerol v/v) using centrifugal filtration (Amicon Ultra-0.5, Millipore Sigma). Protein concentrations were determined using the Qubit Protein Assay Kit (Invitrogen #Q33212). The purity and size of the proteins were validated on a SDS-PAGE gel (Mini-PROTEAN TGX and Mini-TETRA cell, Bio-Rad). Purified proteins were stored at –20° C.

### In vitro transcription (IVT) reactions

Homemade IVT reactions were set up by adding the following components listed at their final concentration: IVT buffer (40 mM Tris-HCl pH 8, 8 mM MgCl_2_, 10 mM DTT, 20 mM NaCl, and 2 mM spermidine), 11.4 mM NTPs pH 7.5, 0.3U thermostable inorganic pyrophosphatase (#M0296S, New England Biolabs), transcription template, DNA gate(s) and MilliQ ultrapure H_2_O to a total volume of 20 μL. Regulated IVT reactions additionally included a purified aTF at the indicated concentration and were equilibrated at 37° C for ~10 minutes. Immediately prior to plate reader measurements, 2 ng of T7 RNAP and, optionally, a ligand at the indicated concentration were added to the reaction. Reactions were then characterized on a plate reader as described in **Plate reader quantification and micromolar equivalent fluorescein (MEF) standardization**.

### RNA extraction from IVT reactions

For RNA products shown on the gel images of **Fig. 2e, Supp. Fig. 3c** and **Supp. Fig. 4c**, IVT reactions were first set up as described above. Then, phenol-chloroform extraction followed by ethanol precipitation was performed to remove any proteins. The reactions were then rehydrated in 1X TURBO™ DNase buffer with 2U of TURBO™ DNase (Invitrogen #QAM2238) to a total volume of 20 μL and incubated at 37° C for 30 minutes to remove the DNA gates and the transcription templates. Then, phenol-chloroform extraction followed by ethanol precipitation was performed again to remove DNase and rehydrated in MilliQ ultrapure H_2_O. The concentrations of the extracted RNA products were measured using the Qubit RNA HS assay kit (Invitrogen #Q32852) and stored in −20° C until further analysis such as PAGE.

### Freeze-drying

Prior to lyophilization, PCR tube caps were punctured with a pin to create three holes. Lyophilization of ROSALIND reactions was then performed by assembling the components of IVT (see above) with the addition of 50 mM sucrose and 250 mM D-mannitol. Assembled reaction tubes were immediately transferred into a pre-chilled aluminum block and placed in a −80° C freezer for 10 minutes to allow slow-freezing. Following the slow-freezing, reaction tubes were wrapped in Kimwipes and aluminum foil, submerged in liquid nitrogen and then transferred to a FreeZone 2.5 L Bench Top Freeze Dry System (Labconco) for overnight freeze-drying with a condenser temperature of −85° C and 0.04 millibar pressure. Unless rehydrated immediately, freeze-dried reactions were packaged as follows. The reactions were placed in a vacuum-sealable bag with a desiccant (Dri-Card Desiccants, Uline #S-19582), purged with Argon using an Argon canister (ArT Wine Preserver, Amazon #8541977939) and immediately vacuum-sealed (KOIOS Vacuum Sealer Machine, Amazon #TVS-2233). The vacuum-sealed bag then was placed in a light-protective bag (Mylar open-ended food bags, Uline #S-11661), heat-sealed (Metronic 8 inch Impulse Bag Sealer, Amazon #8541949845) and stored in a cool, shaded area until usage.

### Plate reader quantification and micromolar equivalent fluorescein (MEF) standardization

A NIST traceable standard (Invitrogen #F36915) was used to convert arbitrary fluorescence measurements to micromolar equivalent fluorescein (MEF). Serial dilutions from a 50 μM stock were prepared in 100 mM sodium borate buffer at pH 9.5, including a 100 mM sodium borate buffer blank (total of 12 samples). The samples were prepared in technical and experimental triplicate (12 samples X 9 replicates = 108 samples total), and fluorescence values were read at an excitation wavelength of 495 nm and emission wavelength of 520 nm for 6-FAM (Fluorescein)-activated fluorescence, or at an excitation wavelength of 472 nm and emission wavelength of 507 nm for 3WJdB-activated fluorescence on a plate reader (Synergy H1, BioTek). Fluorescence values for a fluorescein concentration in which a single replicate saturated the plate reader were excluded from analysis. The remaining replicates (9 per sample) were then averaged at each fluorescein concentration, and the average fluorescence value of the blank was subtracted from all values. Linear regression was then performed for concentrations within the linear range of fluorescence (0–3.125 μM fluorescein) between the measured fluorescence values in arbitrary units and the concentration of fluorescein to identify the conversion factor. For each plate reader, excitation, emission and gain setting, we found a linear conversion factor that was used to correlate arbitrary fluorescence values to MEF (**Supp. Fig. 1, Supplementary Data File 3**).

To characterize reactions, 19 μL of reactions were loaded onto a 384-well optically-clear, flatbottom plate using a multichannel pipette, covered with a plate seal and measured on a plate reader (Synergy H1, BioTek). Kinetic analysis was performed by reading the plate at 1 minute intervals with excitation and emission wavelengths of 495 nm and 520 nm, respectively, for two hours at 37° C. Arbitrary fluorescence values were then converted to MEF by dividing with the appropriate calibration conversion factor.

Besides the data in **Fig. 6b**, no background subtraction was performed when analyzing outputs from any reaction. An example of this standardization procedure is shown in **Supp. Fig. 1**.

### Fluorescence data normalization (Fig. 6b only)

Data shown in **Fig. 6b** were generated by the following normalization method in order to compare experimental observations to ODE simulations. Raw fluorescence values were first standardized to MEF (μM fluorescein) using the method described above. Then, the maximum MEF value was determined among all of the reactions run (5 conditions X 3 replicates = 15 reactions). Each MEF value at every time interval was then normalized using the following formula:

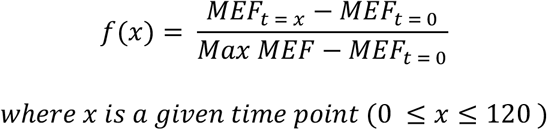

Background subtraction was performed to account for the non-zero fluorescence observed for the quenched DNA signal gate. Once all data were normalized according to the formula above, n=3 replicates per condition were averaged, and the corresponding standard deviation value per condition was calculated.

### Gel Image Analysis

Uncropped, unprocessed gel images presented in **Fig. 2e, Supp. Fig. 2d, Supp, Fig. 3c** and **Supp. Fig 4c** are available as **Supplementary Data File 2** and deposited in Mendeley Data (doi: 10.17632/hr3j3yztxb.1). The band intensity from a SYBR gold stained urea-PAGE gel in **Supp. Fig. 4c** was calculated using Fiji-ImageJ using the traditional lane-profile method as previously described [55]. Briefly, a region of interest in every lane was registered using a rectangle of the same dimension. Then, the uneven background was accounted for by drawing a straight line at the bottom of each peak, and the peak area in each lane was calculated using the wand tool. The peak areas of the RNA standard were then plotted against the total amounts loaded to create the standard curve in **Supp. Fig. 4d** (a linear range: 0.25 – 2 ng). Using the conversion factor from the standard curve, the concentrations of InvadeR variants were estimated from the peak area values obtained from the wand tool.

## Supporting information

DNA Sequences

Gel Images

All source data

Jupyter Notebook Simulations

## STATISTICS AND REPRODUCIBILITY

The number of replicates and types of replicates performed are described in the legend to each figure. Individual data points are shown, and where relevant, the average ± standard deviation is shown; this information is provided in each figure legend. The type of statistical analysis performed in **Fig. 2b, 3c, 4d** and **Supp. Fig. 6** is described in the legend to each figure. Exact p-values along with degrees of freedom computed from the statistical analysis can be found in **Supplementary Data File 3**.

## DATA AVAILABILITY

All data presented in this manuscript are available as supplementary data files. **Supplementary Data Files 2** and **3** are also deposited in Mendeley Data (doi: 10.17632/hr3j3yztxb.1). All plasmids used in this manuscript are available in Addgene with the identifiers 140371, 140374, 140391 and 140395.

## CODE AVAILABILITY

The Jupyter Notebook file with the Python code used in **Fig. 6** and **Fig. 7** is provided as **Supplementary Data File 4**, and the ODE model used in this manuscript is described in the **Supplementary Method**. The Python code is also available in GitHub at https://git.io/Jtlh1.

## AUTHOR CONTRIBUTIONS

Conceptualization, J.K.J., K.K.A. & J.B.L.; Formal analysis, J.K.J. & J.B.L.; Investigation, J.K.J. & K.K.A.; Methodology, J.K.J., K.K.A. & J.B.L.; Visualization, J.K.J. & J.B.L.; Software, J.K.J.; Writing, J.K.J., K.K.A. & J.B.L.; Data curation, J.K.J. & J.B.L.; Funding acquisition, J.B.L.; Validation, J.K.J. & C.M.A.; Project administration, J.K.J. & J.B.L.; Supervision, J.K.J & J.B.L.

## ACKNOWLEDGEMENTS

We thank Andrea Thompson (Northwestern University) and Charlotte Knopp (Northwestern University) for managing the experimental reagents and equipment used in this study; Chloé Archuleta (Northwestern University) for validating the ODE model; Dr. Josiah Hester (Northwestern University), Alex Curtiss (Northwestern University) and John Mamish (Northwestern University) for helpful discussion on the ADC circuit architecture; Dr. Sam Schaffer (National Institute of Standards and Technology) for manuscript editing and helpful discussion on TMSD circuits; Justin Peruzzi (Northwestern University) for assistance with NanoDrop measurements; Dr. Sergii Pshenychnyl (Recombinant Protein Production Core at Northwestern University) for assistance in protein purification. J.K.J. was supported in part by Northwestern University’s Graduate School Cluster in Biotechnology, System, and Synthetic Biology, which is affiliated with the Biotechnology Training Program and by a Ryan Fellowship. This work was also supported by funding from NSF CAREER (1452441 to J.B.L.), NSF MCB RAPID (1929912 to J.B.L.), support from the Crown Family Center for Jewish and Israel Studies at Northwestern University (to J. B. L.), and Searle Funds at The Chicago Community Trust (to J.B.L.).

