## Supplementary Information for "Programming Cell-Free Biosensors with DNA Strand Displacement Circuits"

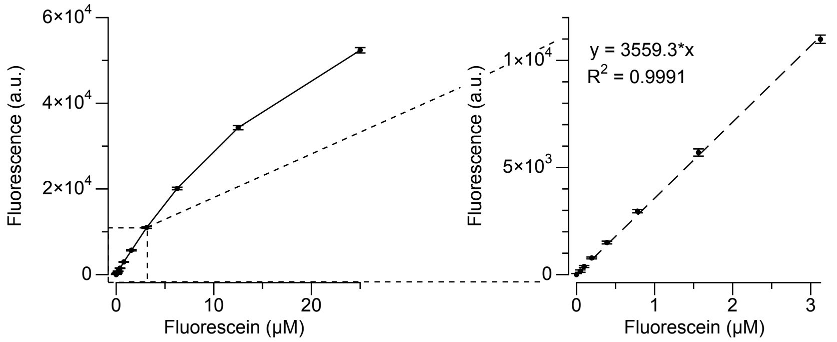


**Supplementary Fig. 1 | Micromolar Equivalent Fluorescein (MEF) standardization**. Arbitrary units of fluorescence were standardized to μM concentrations of fluorescein using a NIST traceable standard (see **Materials and Methods**). In the representative example shown here, a dilution series of fluorescein was prepared in buffer (100 mM sodium borate, pH 9.5) and measured on a plate reader using the same settings for measuring 6' FAM signal (495 nm excitation, 520 nm emission) from the DNA signal gates. The resulting curve, calculated over the linear range of 0–3.125 μM, was then used to standardize fluorescence measured from ROSALIND reactions with TMSD outputs. The standard curve was generated for each plate reader and each measurement setting. Data shown are for n=9 replicates (3 experimentally independent replicates each with 3 technical replicates). Error bars indicate standard deviation computed over n=9 replicates.

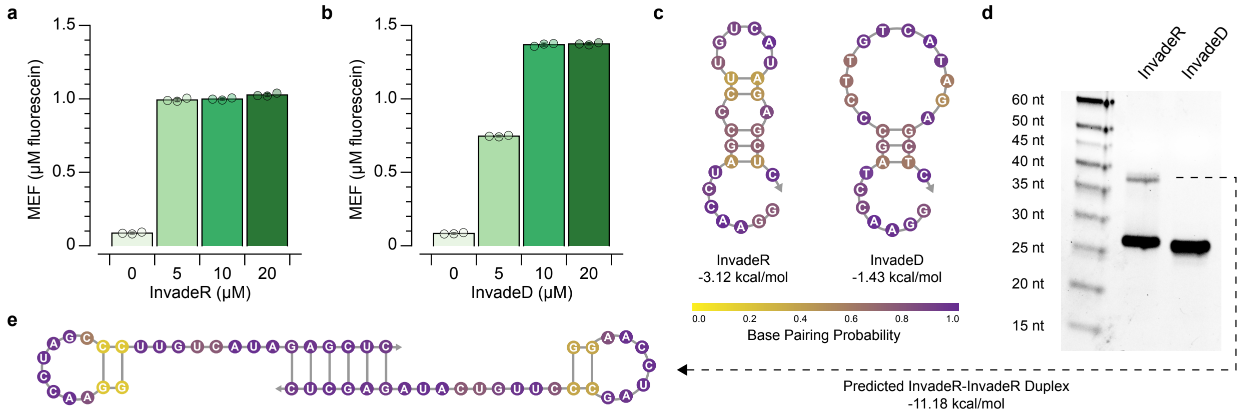


**Supplementary Fig. 2 | TMSD by Invading DNA and RNA strands**. Titration of a purified **a**, InvadeR (RNA) and **b**, InvadeD (DNA) into reactions containing 10 µM of the DNA signal gate in annealing buffer (100 mM potassium acetate, 30 mM HEPES) after 15 minutes. **c**, Secondary structures, minimum free energies and base pairing probabilities of the InvadeR and InvadeD molecules predicted by NUPACK at 37° C [1]. **d**, A urea-PAGE gel of InvadeR and InvadeD. The higher molecular weight band in the InvadeR lane likely corresponds to **e**, a duplex formed by two InvadeR molecules interacting with each other, as predicted by NUPACK. Data shown in **a** and **b** for n=3 technical replicates as points (**a, b**) with bar heights representing the average. Error bars indicate the average value of the replicates ± standard deviation. Data shown for n=1 in **d**. The uncropped, unprocessed gel image shown in **d** is available as **Supplementary Data File 2**.

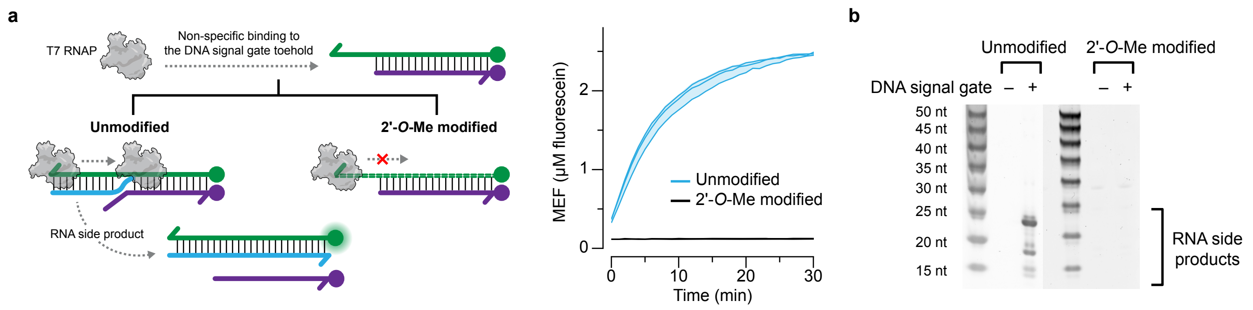


**Supplementary Fig. 3 | Modifying the DNA gate with 2'-*O*-methyl oligonucleotides prevents promoter-independent transcription by T7 RNAP. a**, When the DNA signal gate with its toehold on the 3' end is modified with 2'-*O*-methyl oligonucleotides, no fluorescence activation is observed in the absence of a T7 RNAP-driven IVT template. **b**, When the reactions from **a** were run on a urea-PAGE gel, no RNA side products were observed from the 2'-*O*-methyl DNA signal gate. Data shown in **a** for n=3 technical replicates shown as lines with raw fluorescence standardized to MEF (µM fluorescein). Shading indicates the average value of the replicates ± standard deviation. Data shown for n=1 in **b**. The uncropped, unprocessed gel image shown in **b** is available as **Supplementary Data File 2**.

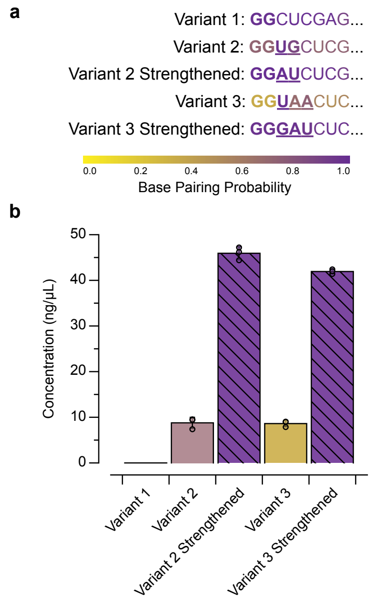


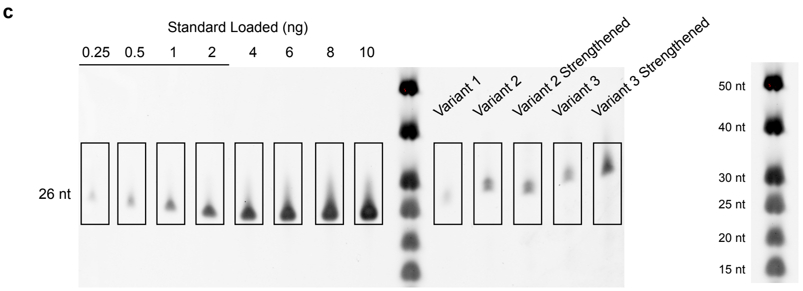


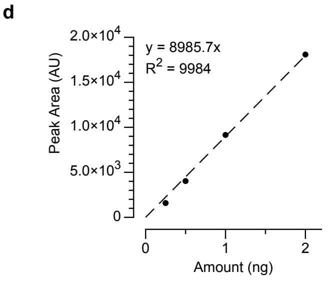


**e**

| | | Amount (ng) | |
|---|---|---|---|
| RNA | Peak Area | PAGE | Qubit |
| Variant 1 | 2640.134 | 0.29 | N/A |
| Variant 2 | 9404.631 | 1.05 | 0.954 |
| Variant 2 Strengthened | 10291.167 | 1.15 | 4.72 |
| Variant 3 | 7431.388 | 0.83 | 0.896 |
| Variant 3 Strengthened | 20183.078 | 2.25 | 4.24 |

**Supplementary Fig. 4 | Transcription efficiency impacts the speed of TMSD. a**, The eight initially transcribed nucleotides of each InvadeR variant in **Fig. 3a**. Nucleotides that are not part of the InvadeR sequences are bolded. **b**, Concentrations of each variant from T7 transcription reactions measured by the Qubit RNA HS assay kit (Invitrogen #Q32852). Each variant was produced *in situ* in the presence of the DNA signal gate for 30 min and extracted (see the RNA extraction from IVT reactions section in **Materials and Methods**). The concentration of variant 1 was too low for Qubit quantification. **c**, The samples measured in **b** were run on a urea-PAGE gel and stained with SYBR gold. Titration of an RNA standard of a similar length was performed to determine the linear range of band peak area quantified by Fiji–ImageJ [2]. **d**, A calibration curve was constructed by plotting the peak area computed from Fiji–ImageJ quantification against the total amount of standard loaded. **e**, Using the calibration curve in **d**, the total amount of RNA for each variant was determined and compared to the measurements made by Qubit in **b**. Data in **b** are shown for n=3 independent experimental replicates as points with bar heights representing the averages. Error bars indicate the average value of the replicates ± standard deviation. Data shown for n=1 in **c**. The uncropped, unprocessed gel image shown in **c** is available as **Supplementary Data File 2**.

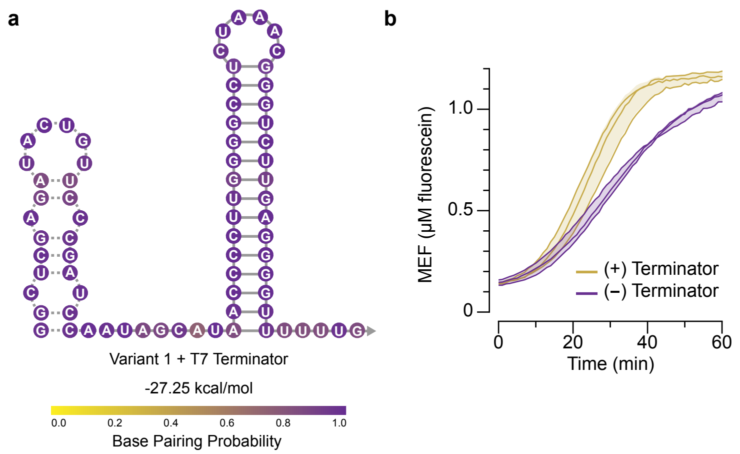


**Supplementary Fig. 5 | Adding a T7 terminator does not notably improve the initial speed of ROSALIND with TMSD. a**, Secondary structure, minimum free energy and base pairing probabilities of the InvadeR variant 1 (dashed backbone) that includes the T7 terminator (solid backbone) as predicted by NUPACK at 37 °C [1]. **b**, Comparison of the kinetics of InvadeR variant 1 TMSD of the DNA signal gate with and without the T7 terminator. All data shown for n=3 independent experimental replicates shown as lines with raw fluorescence standardized to MEF (µM fluorescein). Shading indicates the average value of the replicates ± standard deviation (**Supplementary Data File 3**).

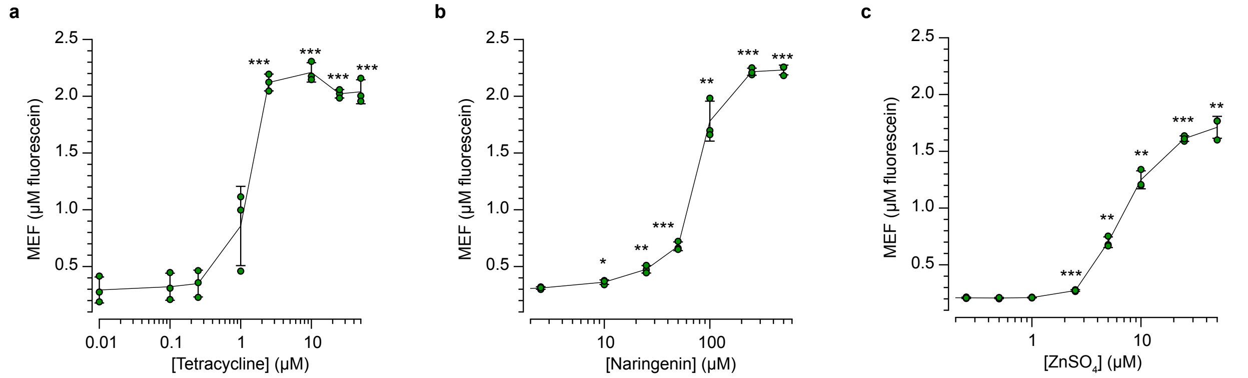


**Supplementary Fig. 6 | Dose response curves of ROSALIND with TMSD.** The dose response curves of ROSALIND with TMSD induced by **a**, tetracycline, **b**, naringenin and **c**, zinc are presented (1h end-point data). The amounts of DNA template and aTF used in each panel are configured as described in **Fig. 5d–f**. All data shown for n=3 independent experimental replicates as points with raw fluorescence values standardized to MEF (μM fluorescein). Error bars indicate the average value of the replicates ± standard deviation. The ligand concentrations at which the signals are distinguishable from the background were determined using a two-tailed, heteroscedastic Student's t-test against the no-ligand condition, and their P value ranges are indicated with asterisks (***P < 0.001, **P = 0.001–0.01, *P = 0.01–0.05). Exact P values along with degrees of freedom can be found in **Supplementary Data File 3**. Data for the no-ligand condition were excluded because the x-axis is on the log scale and are presented in **Supplementary Data File 3**.

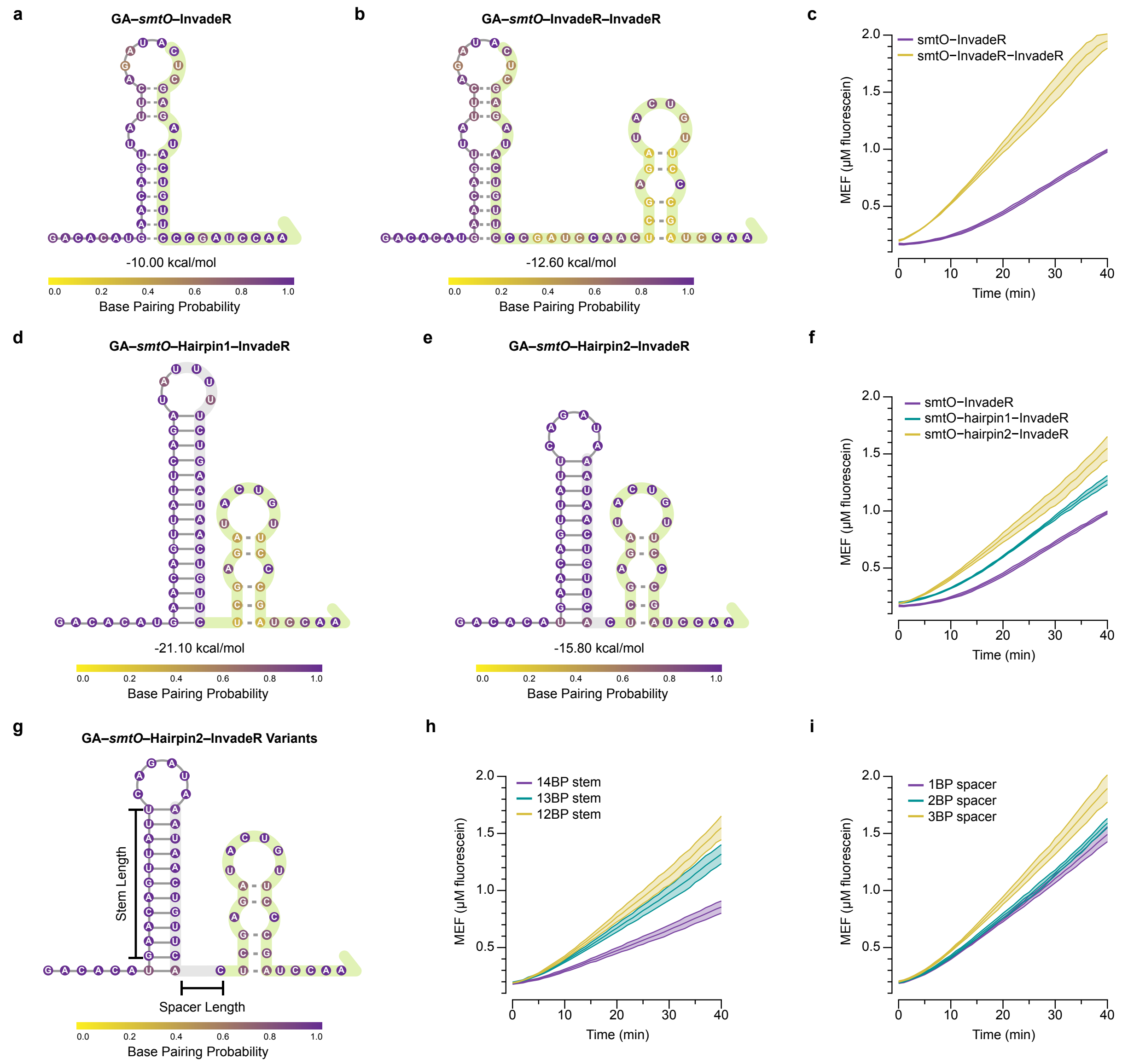


**Supplementary Fig. 7 | Extra nucleotides can be added between the operator and InvadeR sequence to tune the kinetics of the zinc sensor. a**, Secondary structure, minimum free energy and base pairing probabilities of *smtO*-InvadeR predicted by NUPACK at 37° C [1]. A part of the wild type *smtO* sequence is complementary to the InvadeR sequence used, forming a strong predicted stem-loop. The nucleotides highlighted in green correspond to the InvadeR sequence that is designed to strand-displace the DNA signal gate. **b**, Secondary structure, minimum free energy and base pairing probabilities of *smtO*-InvadeR-InvadeR predicted by NUPACK at 37° C. **c**, Comparison of the kinetics of the unregulated *smtO*-InvadeR reaction and the unregulated *smtO*-InvadeR-InvadeR reaction. **d**, **e**, NUPACK-predicted secondary structures, minimum free energies and base pairing probabilities of the two *smtO*-Hairpin-InvadeR variants internally sequester the *smtO* sequence and prevent it from binding to the InvadeR sequence. The nucleotides highlighted in grey indicate the added sequestering sequence. **f**, Comparison of the kinetics of the unregulated reactions of the variants shown in **d** and **e**. **g**, The *smtO*-Hairpin2-InvadeR variants were built by lengthening either the stem length or the spacer between the added hairpin sequence and the InvadeR sequence. The stem length variants are built with the spacer = 0-nt, and the spacer variants are built with the stem length = 12BP. **h**, **i**, Comparison of the kinetics of the unregulated reactions of the *smtO*-Hairpin2-InvadeR variants. All data shown for n=3 independent experimental replicates as lines with raw fluorescence values standardized to MEF (µM fluorescein). Shadings indicate the average value of the replicates ± standard deviation (**Supplementary Data File 3**).

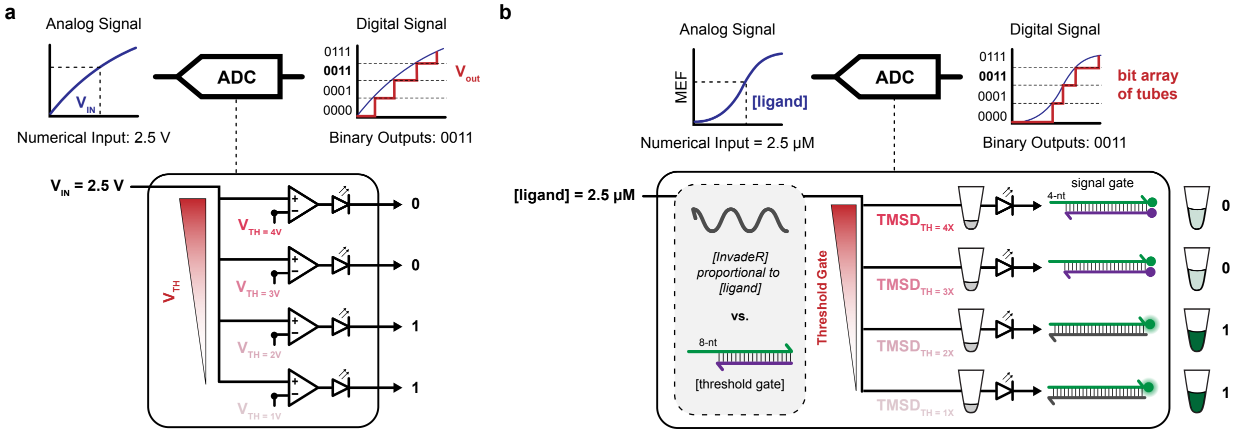


**Supplementary Fig. 8 | A molecular analog-to-digital converter circuit enables ligand quantification. a**, An electronic analog-to-digital converter (ADC) circuit takes in a numerical voltage value (analog input) and generates a binary output of 1s and 0s (digital output). It is built by configuring a series of comparator circuits, each comparing the input voltage to a variable reference voltage to produce a binary output of 1 if the input exceeds the reference value. By setting the threshold voltages ($V_{TH}$) of the comparators to be increasing along the series, the input voltage value is converted into an array of bits with the more 1's representing a higher voltage. **b**, A molecular version of an ADC circuit can be built by implementing TMSD thresholding circuits that compare the input target ligand concentration to a pre-defined threshold value. By titrating the pre-defined threshold value, the molecular ADC circuit can generate different bit arrays to indicate the concentration range of the target ligand.

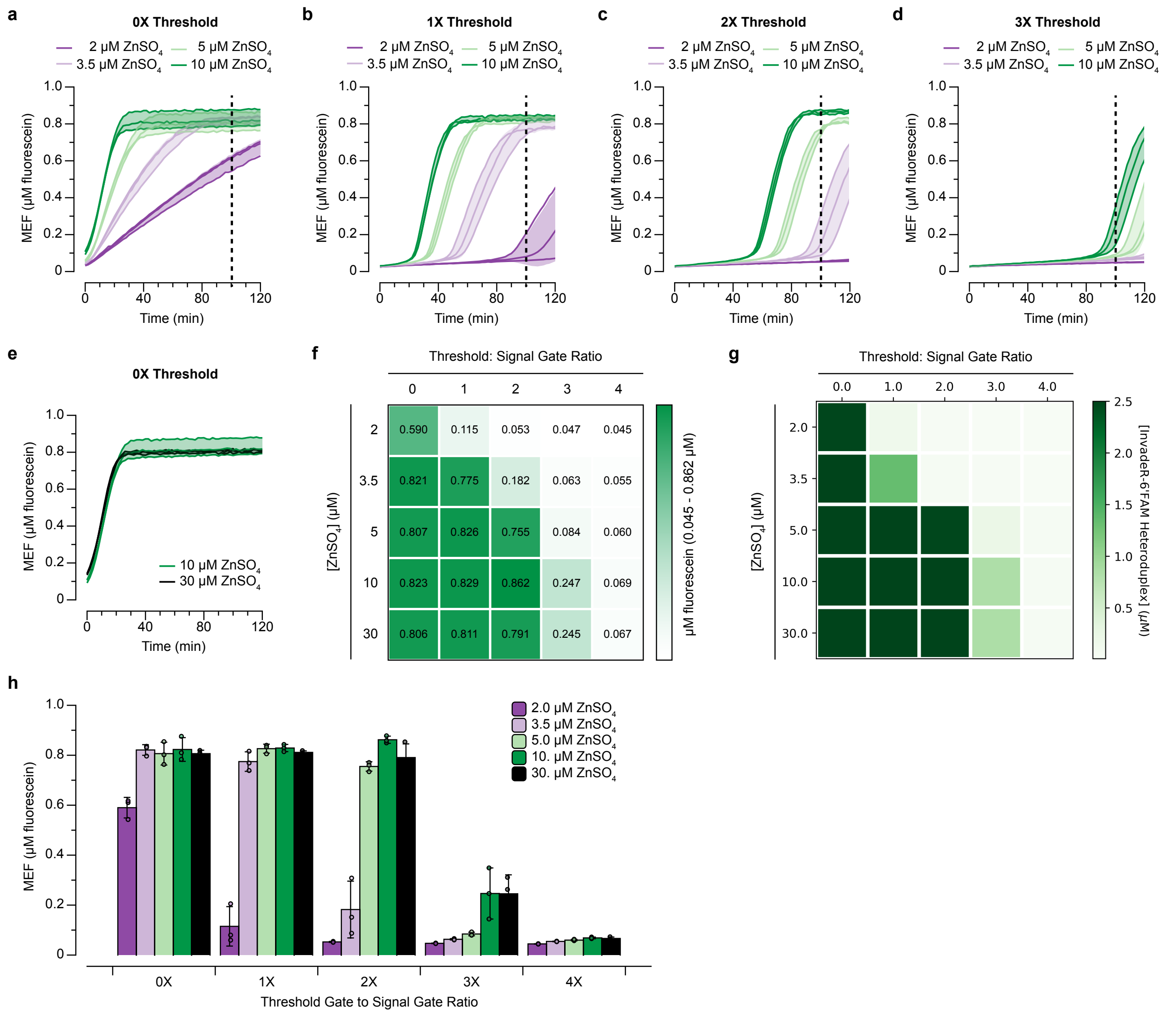


**Supplementary Fig. 9 | The semi-quantitative standard generated by the genetic ADC circuit built with the zinc sensor.** The kinetic traces corresponding to the data shown in **Fig. 6c** are presented for **a**, 0X threshold, **b**, 1X threshold, **c**, 2X threshold and **d**, 3X threshold. The time point (t = 100 min) at which the heatmap was created is indicated with a vertical dotted line. The differences in the response speed for different zinc concentrations are essential in creating the standard. **e**, The 10 μM and 30 μM zinc conditions show no kinetic differences without any threshold gate. **f**, **g**, The functional characterization and ODE predictions of the ADC circuit at 100 min from **Fig. 7c**, **b**, respectively, including the 30 μM zinc and 4X threshold conditions. The 10 μM and 30 μM zinc conditions behave identically as predicted by the ODE model due to their identical kinetic behavior observed in **e**. **h**, The corresponding bar graph data of the semi-quantitative standard shown in **f**. All data shown for n=3 independent experimental replicates as lines (**a**–**e**) or points (**h**) with raw fluorescence values standardized to MEF (μM fluorescein). The values on heatmap in **f** represent averages of the replicates. Shadings in **a**–**e** and error bars in **h** indicate the average value of the replicates ± standard deviation (**Supplementary Data File 3**).

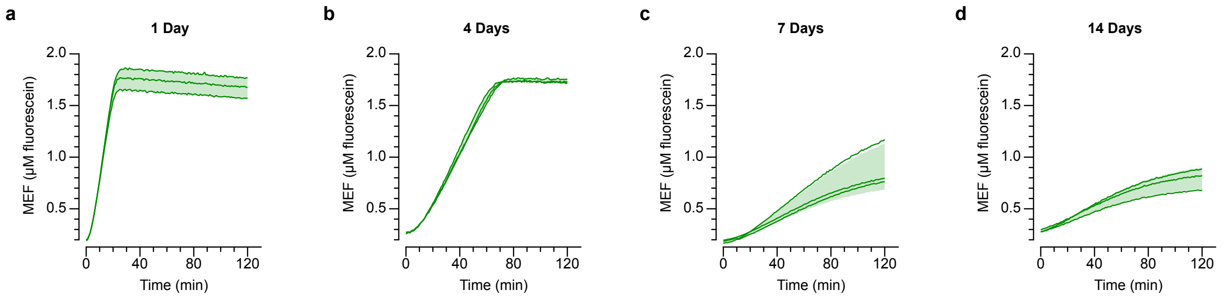


**Supplementary Fig. 10 | ROSALIND with TMSD can be freeze-dried, but improvements are needed for long-term storage.** Unregulated reactions were lyophilized overnight with the addition of 50 mM sucrose and 250 mM D-mannitol as the lyoprotectants. The lyophilized reactions were then vacuum-packaged in a light protective bag with a dri-card and kept in a cool, shaded area until usage. Kinetic traces of rehydrated reactions after **a**, 1 day, **b**, 4 days, **c**, 7 days and **d**, 14 days of storage are shown. There is a decrease in overall signal as well as in the response speed over time. All data shown for n=3 technical replicates as lines with raw fluorescence values standardized to MEF (µM fluorescein). Shadings indicate the average value of the replicates ± standard deviation (**Supplementary Data File 3**).
