## Supplementary Methods for "Programming Cell-Free Biosensors with DNA Strand Displacement Circuits"

### Supplementary Method for Programming Cell-Free Biosensors with DNA Strand Displacement Circuits

#### ODE Model of TMSD Thresholding Circuit

Here, we use the kinetic rates of T7 RNAP-DNA binding, SmtB-*smtO* binding, SmtB-Zn binding, TMSD reactions and T7 RNAP-mediated IVT reactions to simulate ROSALIND Reactions. The following variables will be used:

| Abbreviation | Description |
|---|---|
| D | DNA template |
| RNAP | T7 RNA Polymerase |
| RD | T7 RNAP and DNA template bound complex |
| m | InvadeR |
| TF | Unbound, free SmtB tetramer |
| TFD | SmtB tetramer bound to one *smtO* |
| I | Unbound, free zinc ions |
| TFI | One zinc ion bound to SmtB tetramer |
| RQ | Signal gate |
| SD | InvadeR and 6'FAM heteroduplex |
| Q | Quencher DNA strand |
| Th | Threshold gate |
| SDTh | InvadeR and threshold heteroduplex |
| QTh | Incumbent strand from threshold gate |

In this model, we assume:

1. One-to-one binding of T7 RNAP and T7 promoter on the DNA template
2. The DNA template can be bound to either RNAP or TF, but not both.
3. One-to-one binding of SmtB tetramer and *smtO* on the DNA template (see footnote **a**)
4. One-to-one binding of SmtB tetramer and a zinc ion (see footnote **a**)
5. SmtB tetramer can be bound to either *smtO* on the DNA template or zinc, but not both.

6. All TMSD reactions are irreversible.
7. Fraying within each gate is ignored.

With these assumptions, we have the following reactions and ODEs in the system:

**Reactions:**

$$RNAP + D \underset{k_{unbind}}{\overset{k_{bind}}{\longleftrightarrow}} RD$$

$$TF + D \underset{k_{derepress}}{\overset{k_{repress}}{\longleftrightarrow}} TFD$$

$$TF + I \underset{k_{uninduce}}{\overset{k_{induce}}{\longleftrightarrow}} TFI$$

$$RD \xrightarrow{k_m} m + RNAP + D$$

$$m + RQ \xrightarrow{k_{SD}} SD + Q$$

$$m + Th \xrightarrow{k_{SDTh}} SDTh + QTh$$

**ODEs:**

$$\frac{d[RNAP]}{dt} = k_{unbind}[RD] - k_{bind}[RNAP][D] + k_m[RD]$$

$$\frac{d[RD]}{dt} = k_{bind}[RNAP][D] - k_{unbind}[RD] - k_m[RD]$$

$$\frac{d[D]}{dt} = k_{unbind}[RD] - k_{bind}[RNAP][D] + k_{derepress}[TFD] - k_{repress}[TF][D] + k_m[RD]$$

$$\frac{d[m]}{dt} = k_m[RD] - k_{SD}[RQ][m] - k_{SDTh}[Th][m]$$

$$\frac{d[TF]}{dt} = k_{derepress}[TFD] - k_{repress}[TF][D] - k_{induce}[TF][I] + k_{uninduce}[TFI]$$

$$\frac{d[TFD]}{dt} = k_{repress}[TF][D] - k_{derepress}[TFD]$$

$$\frac{d[I]}{dt} = k_{uninduce}[TFI] - k_{induce}[TF][I]$$

$$\frac{d[TFI]}{dt} = k_{induce}[TF][I] - k_{uninduce}[TFI]$$

$$\frac{d[RQ]}{dt} = -k_{SD}[RQ][m]$$

$$\frac{d[SD]}{dt} = k_{SD}[RQ][m]$$

$$\frac{d[Q]}{dt} = k_{SD}[RQ][m]$$

$$\frac{d[Th]}{dt} = -k_{SDTh}[Th][m]$$

$$\frac{d[SDTh]}{dt} = k_{SDTh}[Th][m]$$

$$\frac{d[QTh]}{dt} = k_{SDTh}[Th][m]$$

This set of ODEs was then run using an ODE solver function, *odeint* from the Scipy.Integrate package in Python 3.7.6. using the rate parameters shown below. For the unregulated reactions shown in Fig. 6b, the initial concentrations of TF, TFD, TFI and I were set to zero.

**Parameters:**

| Parameter | Value | Reference and Note |
|---|---|---|
| $k_m$ for T7-GG-InvR2 | 0.1 /sec | [1] see footnote **b** |
| $k_m$ for T7-GG-smtO-InvR2 | 0.03 /sec | [1] see footnote **b** |
| $k_{bind}$ | 56 /$\mu M$-sec | [2] |
| $k_{unbind}$ | 0.2 /sec | [2] |
| $k_{SD}$ | 0.04 /$\mu M$-sec | [3] for 4-nt toehold |
| $k_{SDTh}$ | 4.0 /$\mu M$-sec | [3] for 8-nt toehold |
| $k_{repress}$ | 3.0 /$\mu M$-sec | see footnote **c** |
| $k_{derepress}$ | 0.18 /sec | see footnote **c** |
| $k_{induce}$ | 80 /$\mu M$-sec | [4] see footnote **d** |
| $k_{uninduce}$ | 0.1 /sec | [4] see footnote **d** |

**Notes:**

a We have found conflicting evidence from literature that reports different DNA-SmtB-zinc binding mechanisms. For instance, *Kar, S. R., et al* reports that SmtB predominantly forms a dimer and binds two zinc ions per subunit (therefore, one SmtB dimer binding 4 total zinc ions) [5]. On the other hand, *VanZile, M. L. et al* reports that SmtB binds one zinc ion per monomer [6]. A more recent literature from *Busenlenher, L. S. et al* reports that two SmtB dimers tightly bind a single 12-2-12 inverted repeat of the *smtO* sequence [7]. We have found that following the mechanism from the most recent literature [7] where SmtB forms a tetramer to bind a single *smtO* site matches our experimental observations the best. We also suspect that while a single SmtB tetramer can bind multiple zinc ions at a time, not all zinc ions are needed to induce the transcription regulated by the SmtB tetramer. Following this logic, we used a model where one zinc ion can bind to the SmtB tetramer to induce the transcription of the invading RNA, which precisely matched our experimental observations.

b These values were adjusted from the reported rate to accommodate for different initially transcribed nucleotides in each template. For instance, pT7-GGGA has a much greater transcription efficiency than pT7-GGCA [8].

c There was no literature with this information available to the best of our knowledge. We have decided to estimate these values based on a reasonable range of a $K_d$ value for a transcriptional repressor, which is typically in the nanomolar range. Currently, the rate constants are set so that the $K_d = \frac{k_{derepress}}{k_{repress}} \approx 60nM$.

d These values have been estimated to fit the association constants of SmtB-Zinc reported in *VanZile, et al* [4].
