## Supplementary material for "Programming Cell-Free Biosensors with DNA Strand Displacement Circuits": Gel Images: Fig.2e_Notes.txt

"Programming Cell-Free Biosensors with DNA Strand Displacement Circuits."
Jung, et al. (2021)

Fig.2e.tif shows the uncropped, unprocessed urea-PAGE gel image of data shown in Fig.2e. Lanes 4, 5, 6, 10 and 11 are not shown in the manuscript. 

From left to right:
 1. Low Molecular ssDNA Ladder (10, 15, 20, 25, 30, 35, 40, 45, 50, 60, 70, 80, 90, 100 nt)
 2. (-) 3' toehold DNA signal gate (-) DNA template
 3. (+) 3' toehold DNA signal gate (-) DNA template

 7. Low Molecular ssDNA Ladder (10, 15, 20, 25, 30, 35, 40, 45, 50, 60, 70, 80, 90, 100 nt)
 8. (-) 5' toehold DNA signal gate (-) DNA template
 9. (+) 5' toehold DNA signal gate (-) DNA template
