## Supplementary material for "Programming Cell-Free Biosensors with DNA Strand Displacement Circuits": Gel Images: Supp.Fig.2d_Notes.txt

"Programming Cell-Free Biosensors with DNA Strand Displacement Circuits."
Jung, et al. (2021)

Supp.Fig.2d.tif shows the uncropped, unprocessed urea-PAGE gel image of data shown in Supp.Fig.2d. Lanes 4 - 10 are not shown in the manuscript. 

From left to right:
1. Low Molecular ssDNA Ladder (10, 15, 20, 25, 30, 35, 40, 45, 50, 60, 70, 80, 90, 100 nt)
2. PAGE-purified InvadeR designed to strand-displace the 3' toehold DNA signal gate
3. PAGE-purified InvadeD designed to strand-displace the 3' toehold DNA signal gate
