## Supplementary material for "Programming Cell-Free Biosensors with DNA Strand Displacement Circuits": Gel Images: Supp.Fig.3b_Notes.txt

"Programming Cell-Free Biosensors with DNA Strand Displacement Circuits."
Jung, et al. (2021)

Supp.Fig.3b.tif shows the uncropped, unprocessed urea-PAGE gel image of 2'-O-methylation modification data shown in Supp.Fig.3b. Lanes 1-6, 10 and 11 are not shown in the manuscript. The 3' toehold DNA signal gate without 2'-O-methylation modification data shown are the same gel image used in Fig.2e.

From left to right:
 6. Low Molecular ssDNA Ladder (10, 15, 20, 25, 30, 35, 40, 45, 50, 60, 70, 80, 90, 100 nt)
 7. (-) 3' toehold DNA signal gate with 2'O-methylation modification (-) DNA template
 8. (+) 3' toehold DNA signal gate with 2'O-methylation modification (-) DNA template
