## Supplementary material for "Programming Cell-Free Biosensors with DNA Strand Displacement Circuits": Gel Images: Supp.Fig.4c_Notes.txt

"Programming Cell-Free Biosensors with DNA Strand Displacement Circuits."
Jung, et al. (2021)

Supp.Fig.4c.tif shows the uncropped, unprocessed urea-PAGE gel image of data shown in Supp.Fig.4c.

From left to right:
 1. 0.25 ng of Standard RNA
 2. 0.5 ng of Standard RNA
 3. 1 ng of Standard RNA
 4. 2 ng of Standard RNA
 5. 4 ng of Standard RNA
 6. 6 ng of Standard RNA
 7. 8 ng of Standard RNA
 8. 10 ng of Standard RNA
 9. Low Molecular ssDNA Ladder (10, 15, 20, 25, 30, 40, 50, 60 nt)
10. InvadeR Variant 1 designed to strand-displace the 5' toehold DNA signal gate extracted from a 30-min long IVT reaction
11. InvadeR Variant 2 designed to strand-displace the 5' toehold DNA signal gate extracted from a 30-min long IVT reaction
12. InvadeR Variant 2 Strengthened designed to strand-displace the 5' toehold DNA signal gate extracted from a 30-min long IVT reaction
13. InvadeR Variant 3 designed to strand-displace the 5' toehold DNA signal gate extracted from a 30-min long IVT reaction
14. InvadeR Variant 3 Strengthened designed to strand-displace the 5' toehold DNA signal gate extracted from a 30-min long IVT reaction
