## Supplementary material for "Programming Cell-Free Biosensors with DNA Strand Displacement Circuits": Jupyter Notebook Simulations: Jung_TMSD_ROSALIND_Supp_Data_File4_ODE_Simulations.ipynb

```
# %% [markdown]
### Modeling TMSD Thresholding Circuit:


Using the kinetics rates of RNAP-DNA binding, SmtB-smtO binding, SmtB-zinc binding, and toehold-mediated strand displacement reactions, simulate ROSALIND Reactions. Following ODEs will be computed:<br>

D - DNA template <br>
RNAP - T7 RNAP <br>
RD - T7 RNAP and DNA template bound complex <br>
m - InvadeR RNA <br><br>
TF - Unbound, free SmtB tetramer <br>
TFD - SmtB tetramer bound to one smtO <br>
I - Unbound, free zinc ion <br>
TFI - One zinc ion bound to SmtB tetramer <br><br>
RQ - Signal gate <br>
SD - InvadeR and 6'FAM heteroduplex <br>
Q - Quencher DNA strand <br>
Th - Threshold gate <br>
SDTh - InvadeR and threshold heteroduplex <br>
QTh - Incumbent strand from threshold gate <br><br><br>


**Reactions:**<br><br>

$$RNAP + D \overset{k_{bind}}{\underset{k_{unbind}}\rightleftarrows} RD$$
<br>
<br>
$$TF + D \overset{k_{repress}}{\underset{k_{derepress}}\rightleftarrows} TFD$$
<br>
<br>
$$TF + I \overset{k_{induce}}{\underset{k_{uninduce}}\rightleftarrows} TFI$$
<br>
<br>
$$RD \xrightarrow[\text{}]{\text{k_m}} m + RNAP + D$$
<br>
<br>
$$m + RQ \xrightarrow[\text{}]{\text{k_SD}} SD + Q$$
<br>
<br>
$$m + Th \xrightarrow[\text{}]{\text{k_SDTh}} SDTh + QTh$$
<br>
<br>

**ODEs:**<br>

$$\dfrac{d[RNAP]}{dt} = k_{unbind}[RD] - k_{bind}[RNAP][D] + k_{m}[RD]$$
<br>
$$\dfrac{d[RD]}{dt} = k_{bind}[RNAP][D] - k_{unbind}[RD] - k_{m}[RD]$$
<br>
$$\dfrac{d[D]}{dt} = k_{unbind}[RD] - k_{bind}[RNAP][D] + k_{derepress}[TFD] - k_{repress}[TF][D] + k_{m}[RD]$$
<br>
$$\dfrac{d[m]}{dt} = k_{m}[RD] - k_{SD}[RQ][m] - k_{SDTh}[Th][m]$$
<br>
<br>
$$\dfrac{d[TF]}{dt} = k_{derepress}[TFD] - k_{repress}[TF][D] - k_{induce}[TF][I] + k_{uninduce}[TFI]$$
<br>
$$\dfrac{d[TFD]}{dt} = k_{repress}[TF][D] - k_{derepress}[TFD]$$
<br>
$$\dfrac{d[I]}{dt} = k_{uninduce}[TFI] - k_{induce}[TF][I]$$
<br>
$$\dfrac{d[TFI]}{dt} = k_{induce}[TF][I] - k_{uninduce}[TFI]$$
<br>
<br>
$$\dfrac{d[RQ]}{dt} = -k_{SD}[RQ][m]$$
<br>
$$\dfrac{d[SD]}{dt} = k_{SD}[RQ][m]$$
<br>
$$\dfrac{d[Q]}{dt} = k_{SD}[RQ][m] $$
<br>
$$\dfrac{d[Th]}{dt} = -k_{SDTh}[Th][m]$$
<br>
$$\dfrac{d[SDTh]}{dt} = k_{SDTh}[Th][m]$$
<br>
$$\dfrac{d[QTh]}{dt} = k_{SDTh}[Th][m]$$
<br>
<br>

**Parameters:**<br>

$$k_{m} = 0.1/\text{sec [1]}$$
<br>
$$k_{bind} = 56 /\mu M\text{-sec} \qquad k_{unbind} = 0.2 \text{/sec [2]}$$
<br>
$$k_{SD} = 0.04 /\mu M\text{-sec}\qquad \text{4 nt toehold [3]}$$
<br>
$$k_{SDTh} = 4.0 /\mu M\text{-sec}\qquad \text{8 nt toehold [3]}$$
<br>
$$k_{repress} = 3.0 /\mu M\text{-sec} \qquad k_{derepress} = 0.18 \text{/sec}$$
<br>
$$k_{induce} = 80 /\mu M\text{-sec} \qquad k_{uninduce} = 0.1 \text{/sec [4]}$$
<br><br>

**References:**<br>
[1] McClure, W.R., *et al.* "Rate-limiting steps in RNA chain initiation." *PNAS.* (1980).<br>

[2] Ujvari, A., *et al.* "Thermodynamics and kinetic measurements of promoter biding by T7 RNA polymerase." *Biochemistry.* (1996).<br>

[3] Srinivas, N., *et al.* “On the biophysics and kinetics of toehold-mediated DNA strand displacement.” *Nucleic Acids Research.* (2013). <br>

[4] VanZile, M. L., *et al.* "Structural Characterization of Distinct a3N and a5 Metal Sites in the Cyanobacterial Zinc Sensor SmtB." *Biochemistry.* (2002).<br>

# %% [code]
# Importing packages:
import numpy as np
import scipy.integrate as spi
import matplotlib.pyplot as plt
import math as mt
import seaborn as sns

# Setting the time range to run the simulation for:
t_start = 0
t_step = 2.
t_stop = 121.
t = np.arange(t_start,t_stop,t_step)

# %% [code]
# Write out the set of ODEs to solve:
def dy(y,t,params):
    [D, RNAP, RD, TF, TFD, I, TFI, m, SD, SDTh, Q, Qth, RQ, Th] = y
    k_m, k_bind, k_unbind, k_SD, k_SDTh, k_repress, k_derepress, k_induce, k_uninduce = params
    
    dD = k_unbind*RD - k_bind*D*RNAP + k_derepress*TFD - k_repress*D*TF + k_m*RD #Free DNA
    dRNAP = k_unbind*RD - k_bind*D*RNAP + k_m*RD #Free RNA polymerase
    dRD = k_bind*D*RNAP - k_unbind*RD - k_m*RD #DNA-RNAP bound complex
    dm = k_m*RD - k_SD*RQ*m - k_SDTh*Th*m #InvadeR
    
    dTF = k_derepress*TFD - k_repress*D*TF - k_induce*TF*I + k_uninduce*TFI #Free SmtB tetramer
    dTFD = k_repress*D*TF - k_derepress*TFD #SmtB-DNA bound complex
    dI = k_uninduce*TFI - k_induce*TF*I #Free Zinc ions
    dTFI = k_induce*TF*I - k_uninduce*TFI #SmtB-Zinc bound complex
    
    dRQ = -k_SD*RQ*m #Signal gate
    dSD = k_SD*RQ*m #InvadeR-Reporter heteroduplex
    dQ = k_SD*RQ*m #Free quencher DNA strand
    
    dTh = -k_SDTh*Th*m #Threshold gate
    dSDTh = k_SDTh*Th*m #InvadeR-Threshold heteroduplex
    dQth = k_SDTh*Th*m #Free threshold incumbent DNA strand
    
    diff = [dD, dRNAP, dRD, dTF, dTFD, dI, dTFI, dm, dSD, dSDTh, dQ, dQth, dRQ, dTh]
    
    return diff

# %% [code]
# Setting the parameters:
k_m = 0.1*60. #Transcription rate of gg-InvadeR_v2 per min
k_bind = 56.*60. #DNA and T7 binding per µM per min
k_unbind = 0.20*60. #DNA and T7 unbinding per min
k_SD = 0.04*60. #SD rate per µM per min 
k_SDTh = 4.*60. #Threshold SD rate per µM per min
k_repress = 3.*60. #DNA-SmtB binding per µM per min
k_derepress = 0.18*60. #DNA-SmtB dissocating per min
k_induce = 80.*60. #SmtB-Zinc binding per µM per min
k_uninduce = 0.1*60. #SmtB-Zinc dissociating per min

# Setting parameters to feed into ODE solver:
params = [k_m, k_bind, k_unbind, k_SD, k_SDTh, k_repress, k_derepress, k_induce, k_uninduce]

# %% [code]
# Setting initial input values in µM:
D_0 = 0.05 #free DNA
RNAP_0 = 0.1 #RNAP
RD_0 = 0.0 #DNA bound to RNAP
m_0 = 0.0 #gg-InvadeR_v2

TF_0 = 0.0 #SmtB tetramer
TFD_0 = 0.0 #SmtB-DNA bound complex
I_0 = 0.0 #Free Zinc ions
TFI_0 = 0.0 #SmtB-Zinc bound complex

RQ_0 = 5.0 #Output gate
SD_0 = 0.0 #InvadeR-Reporter heteroduplex
Q_0 = 0.0 #Quencher ssDNA

Th_0 = [0.0, 5.0, 10., 20., 40.] #Threshold gate
SDTh_0 = 0.0 #InvadeR-Threshold heteroduplex
Qth_0 = 0.0 #Threshold incumbent ssDNA

# Solving for the kinetics per [threshold gate]:
for i in Th_0:
    y_0 = [D_0, RNAP_0, RD_0, TF_0, TFD_0, I_0, TFI_0, m_0, SD_0, SDTh_0, Q_0, Qth_0, RQ_0, i]
        
    # Running the ODEs for each parameter:
    y = spi.odeint(dy,y_0,t,args=(params,))
        
    # Obtaining the variables to plot:
    temp = y[:,8]
    
    # Getting the threshold to signal gate ratio:
    ratio = i/RQ_0 
    
    # Plotting the result:
    plt.plot(t, temp, label=str(ratio)+'X')
    plt.xlabel("Time (min)")
    plt.ylabel("[6'FAM] ($\mu$M)")
    plt.legend(loc='best')

# %% [code]
# Setting the parameters:
k_m = 0.03*60. #Transcription rate of gg-smtO-InvadeR_v2 per min

'''k_m values is lower here than gg-InvadeR_v2, because of the smtO sequence.
   pT7-gg-smtO-InvadeR_v2 ends up being TAATACGACTCACTATAGGC... which has lower Txn efficiecny than
   pT7-gg-InvadeR_v2, which has TAATACGACTCACTATAGGG...'''

# Setting parameters to feed into ODE solver:
params = [k_m, k_bind, k_unbind, k_SD, k_SDTh, k_repress, k_derepress, k_induce, k_uninduce]

# Setting initial input values in µM:
m_0 = 0.0 #gg-smtO-InvadeR_v2

TF_0 = 5. #SmtB tetramer
TFD_0 = 0.0 #SmtB-DNA bound complex
I_0 = [2., 3.5, 5., 10.] #Free Zinc ions
TFI_0 = 0.0 #SmtB-Zinc bound complex

RQ_0 = 2.5 #Output gate
SD_0 = 0.0 #InvadeR-Reporter heteroduplex
Q_0 = 0.0 #Quencher ssDNA

Th_0 = [0.0, 2.5, 5.0, 7.5] #Threshold gate
SDTh_0 = 0.0 #InvadeR-Threshold heteroduplex
Qth_0 = 0.0 #Threshold incumbent ssDNA

ratio = [i/RQ_0 for i in Th_0]
matrix = []

# Solving for the kinetics per [threshold gate]:
for i in range(len(I_0)):
    
    # Create an empty list to fill in the endpoint values at each [ZnSO4] per [threshold gate]:
    temp = []
    
    # Solving for the kinetics per [ZnSO4]
    for j in Th_0:
        y_0 = [D_0, RNAP_0, RD_0, TF_0, TFD_0, I_0[i], TFI_0, m_0, SD_0, SDTh_0, Q_0, Qth_0, RQ_0, j]
        
        # Running the ODEs for each parameter:
        y = spi.odeint(dy,y_0,t,args=(params,))
        
        # Obtaining the endpoint to plot:
        temp.append(y[:,8][50])
        
    matrix.append(temp)

# Plotting the result:
matrix = np.array(matrix)

fig = plt.figure()
fig, ax = plt.subplots(dpi=500)
heatmap = ax.imshow(matrix, cmap = 'Greens')

# Setting and labelling axis:
ax.set_xticks(np.arange(len(ratio)))
ax.set_yticks(np.arange(len(I_0)))
ax.set_xticklabels(ratio)
ax.set_yticklabels(I_0)
ax.set_xlabel("Threshold to Output Gate Ratio\n")
ax.set_ylabel("[ZnSO$_4$] ($\mu$M)\n")
ax.xaxis.set_label_position('top')
ax.tick_params(top=True, bottom=False, labeltop=True, labelbottom=False)

# Creating a colorbar:
cbar = ax.figure.colorbar(heatmap)
cbar.ax.set_ylabel("[InvadeR-6'FAM Heteroduplex] ($\mu$M)", rotation=-90, va="bottom")

# Creating the white lines between each cell:
ax.set_xticks(np.arange(matrix.shape[1]+1)-.5, minor=True)
ax.set_yticks(np.arange(matrix.shape[0]+1)-.5, minor=True)
ax.grid(which="minor", color="w", linestyle='-', linewidth=3)
ax.tick_params(which="minor", bottom=False, left=False)

# Saving the image:
plt.savefig('Zinc_Threshold_Heatmap_4X4', bbox_inches="tight")

# %% [code]


# %% [code]
```
